# Parallel circuits in the posterior parietal cortex balance behavioral flexibility and stability

**DOI:** 10.1101/2025.06.29.662178

**Authors:** Eunji Jung, Jong-Hoon Lee, Woochul Choi, Gil-Hyun Kim, Geonhui Ryu, Se-Bum Paik, Seung-Hee Lee

## Abstract

Rapid behavioral adaptation requires the brain to solve a fundamental computational dilemma: how to flexibly update learned rules while maintaining stable motor performance. This flexibility-stability trade-off is central to cognitive function and artificial intelligence, yet its neural circuit basis is poorly understood. Here, we identify the circuit architecture that coordinates this balance. Using a within-session auditory reversal learning task combined with circuit-specific manipulations in mice, we demonstrate that the posterior parietal cortex (PPC) orchestrates adaptive learning through two parallel, anatomically distinct pathways to the auditory cortex (AC) and inferior colliculus (IC). These two circuits perform a double dissociation of function: the PPC-to-AC circuit facilitates flexible rule updating by encoding dynamic stimulus-outcome contingencies, while the PPC-to-IC circuit preserves stable action execution across changing rules. A feedforward network model further demonstrates that segregated stable and flexible top-down projections are necessary and sufficient for rapid reversal learning. This parallel architecture segregates adaptive and stable computations into distinct circuits, providing a mechanistic framework for cognitive flexibility and offering potential insight into conditions where this balance is disrupted, such as obsessive-compulsive disorder and schizophrenia.

## Introduction

Behavioral flexibility, the ability to adapt behavioral responses to changes in environmental contexts even under identical sensory conditions, requires flexible association of sensory information with appropriate motor outputs^1,2^. To achieve rapid adaptation to rule changes, the brain must maintain stable, goal-directed actions while simultaneously updating task rules to support successful behavioral transitions according to new rules^3,4^ Disruption of this coordination is a hallmark of several neurological and psychiatric disorders. For instance, patients with schizophrenia often exhibit unconstrained action patterns and impaired stability in goal-directed behaviors^5,6^, whereas those with Parkinson’s disease or obsessive-compulsive disorder demonstrate maladaptation to new rules, characterized by excessive perseveration^2,7–10^. This flexibility-stability dilemma also poses a significant challenge in computational learning, and modern machine-learning algorithms address this by balancing flexible updates with stable maintenance through functional modularization^11–13^. However, the neural mechanisms by which the brain negotiates this flexibility-stability trade-off during adaptive learning, enabling the acquisition of new behaviors without compromising execution stability, remain poorly understood. It is still debated whether this balance is maintained within a single network through specific representational dynamics or whether it relies on anatomically segregated networks operating in parallel.

Reversal learning tasks, which require updating behavioral responses following reversed stimulus-outcome contingencies, provide a powerful paradigm for interrogating behavioral flexibility^2,14^. While the prefrontal cortex has traditionally been considered central to behavioral flexibility through the encoding of task rules^15–17^, recent evidence suggests that a broader network of brain areas dynamically adjusts representations in response to task-relevant contingencies^18–20^. The posterior parietal cortex (PPC), with its extensive bidirectional connections to both sensory and frontal areas^21,22^, has emerged as a critical node in this process. In flexible learning tasks, the PPC encodes task-relevant choices rather than raw sensory stimuli^23^, is required for rule-based sensory categorization^24^, and retains a history of behavioral contingencies to guide future decisions^25,26^. These properties suggest that the PPC acts as a flexible hub that can update outcome contingencies during reversal learning, potentially through adaptive control of inter-areal connectivity and compositional coding^27,28^. However, the underlying circuit architecture that implements these principles while simultaneously ensuring behavioral stability during rapid transitions is a fundamental question that has yet to be addressed.

During reversal learning, the frontal cortex exhibits increased activity associated with rule updating^29,30^, while sensory cortical activity is enhanced only after full adaptation to new contingencies^31^. While sensory representations are known to be modulated to prioritize task-relevant stimuli^32–34^, how stability and flexibility are manifested across distinct sensory areas during adaptive perceptual performance has yet to be fully elucidated. Recent evidence indicates that both sensory and context-dependent prior information is distributed across multiple brain areas^35,36^. Remaining question is how such priors shape sensory representations across the sensory hierarchy and, more importantly, whether these representational dynamics are causally linked to flexible behavioral outcomes. In the auditory system, both the midbrain inferior colliculus (IC) and the auditory cortex (AC) exhibit significant response modulations driven by task demands^34,37,38^ and internal behavioral states^39–42^. Furthermore, both structures encode non-sensory information, including action-and reward-related signals^38,43–49^, suggesting that top-down influences orchestrate auditory processing across the auditory pathway. Nevertheless, a systematic comparison of cortical and subcortical auditory processing during the course of behavioral adaptation and how they differentially contribute to the stability and flexibility of auditory behavior is currently lacking.

In this study, we demonstrate that parallel top-down projections from the PPC to the AC and IC enable rapid behavioral adaptation during within-session auditory Go/No-go reversal learning. We first found that AC neurons flexibly alter their representations during reversal learning to enhance cue discrimination, whereas IC neurons maintain stable representations across rules. Consistent with these observations, pharmacological inactivation of the AC impaired adaptation to reversed contingencies, while silencing the IC disrupted auditory discriminability regardless of contingency. Using dual retrograde tracing, we identified anatomically segregated PPC subpopulations projecting to the AC (PPC_AC_) and IC (PPC_IC_). Projection-specific calcium imaging and optogenetic perturbations revealed that PPC_AC_ neurons flexibly update stimulus-outcome associations, while PPC_IC_ neurons support stable, goal-directed action execution during reversal learning. Finally, simulations using a feed-forward network model indicated that this parallel architecture is a computational necessity for rapid adaptation following contingency reversal. Together, our findings reveal a circuit-level division of labor within the PPC that balances flexibility and stability to support rapid and robust behavioral adaptation.

## Result

### Fast transitions of behavioral responses in mice performing a within-session auditory reversal learning

To examine behavioral flexibility during rapid contingency changes, we developed a within-session auditory reversal-learning task for head-fixed mice using a Go/No-go paradigm (Figure 1A; Methods). Mice were initially trained under the first rule (R1) to lick in response to a low-frequency tone (Go, 5kHz) and to withhold licking in response to a high-frequency tone (No-go, 10kHz). Upon reaching expert-level performance (correct response rate > 0.75 within 200 trials), stimulus-outcome contingencies were reversed within the same session (R2; Go, 10 kHz; No-go, 5 kHz).

**Figure 1.**
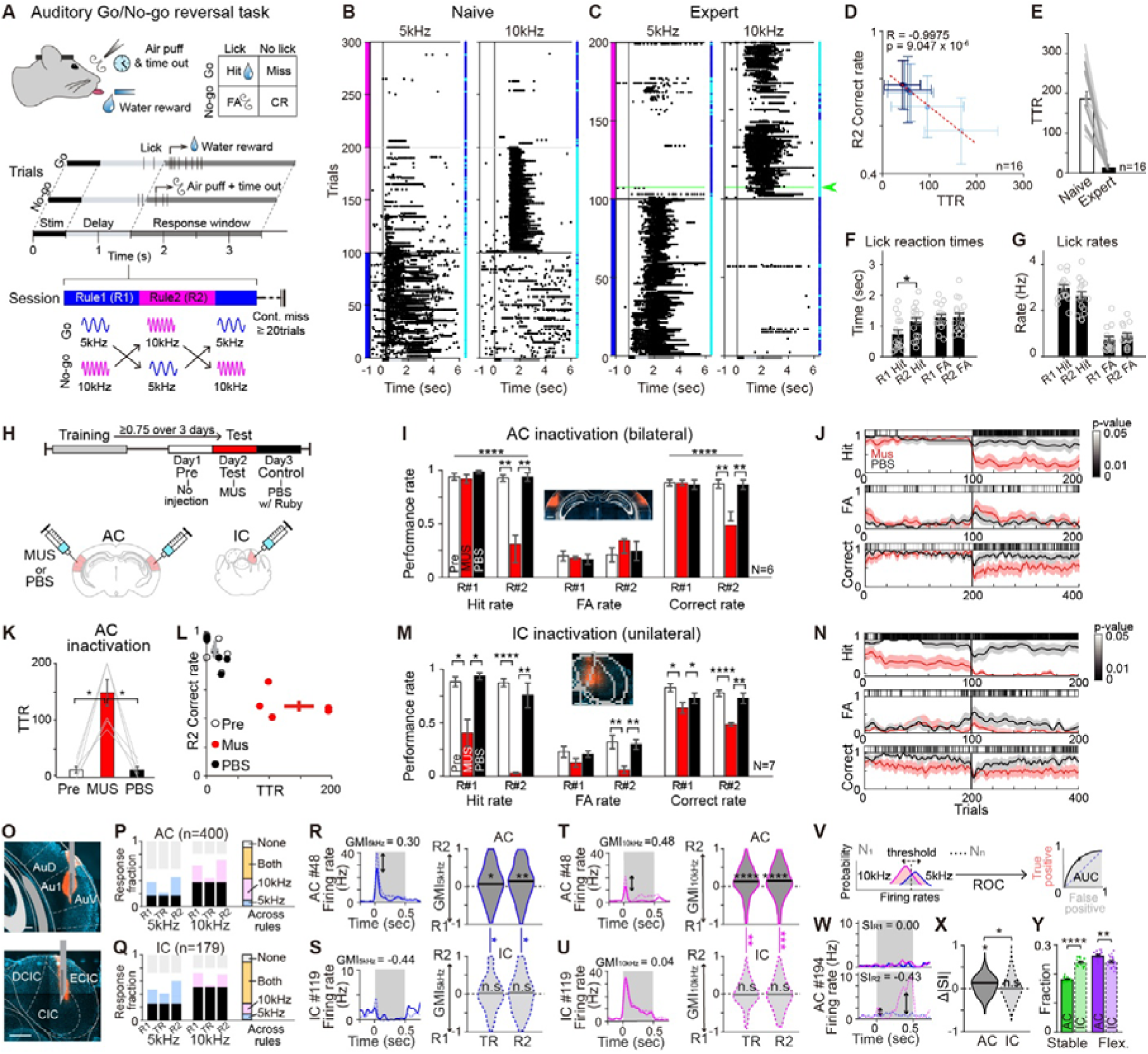
Distinct functional roles of AC and IC in rapid reversal learning. **A.** Schematic of the auditory Go/No-go reversal-learning task. **B-C.** Example lick raster plots from a naïve (**B**) and expert (**C**) mouse during reversal learning. Left, 5kHz trials; right, 10kHz trials. Each black dot represents a single lick. Blue and magenta vertical bands denote R1 and R2 blocks, respectively. Vertical black lines denote sound onset (0 s); horizontal black lines indicate the rule-reversal point; green line and arrow mark the TTRs; cyan dots indicate lick trials; blue dots indicate no-lick trials. Naïve mice exhibited licking after passive water delivery during R2 (**B**), whereas expert mice initiated licking in response to the 10kHz stimulus following reversal (**C**). **D.** Scatter plot of TTR vs. R2 correct rates across training sessions. Sessions for each mouse were divided into six equal training segments, averaged within each segment, and then averaged across mice (one circle per segment). Red dashed line indicates linear regression. Pearson’s correlation shows improved performance with training. **E.** Average TTRs for naïve (189.82 ± 19.04) and expert (30.82 ± 6.25) mice, demonstrating faster adaptation after training. **F.** Lick reaction times for hit and FA trials. Hit reaction times were significantly longer in R2 than in R1 (p = 0.0275), whereas FA reaction times did not differ. Wilcoxon rank-sum test. **G.** Lick rates for hit and FA trials, showing no significant differences across rules. **H.** Schematic of MUS or PBS injections into bilateral AC or unilateral IC. **I-L.** Behavioral effects of AC inactivation (N=6). **I.** Average hit, FA, and correct rates during Pre (no injection; open bars), MUS (red bars), and PBS (black bars) sessions. Repeated measures two-way ANOVA; hit rate (drug, p = 7.88 x 10^-7^; rule, p = 3.53 x 10^-5^; interaction, p = 6.69 x 10^-6^); FA rate (drug, p = 0.566; rule, p = 0.409; interaction, p = 0.850); correct rate (drug, p = 1.04 x 10^-5^; rule, p = 9.3 x 10^-5^; interaction, p = 4.83 x 10^-6^). Inset shows bilateral AC injection sites aligned to the Paxinos atlas. Scale bar, 500 μm. **J.** Session-averaged performance with AC inactivation. Hit, FA, and correct rates are shown for MUS (red) and PBS (black). Shading indicates ± SEM. Grayscale bars denote p-values for significant differences between MUS and PBS (Wilcoxon signed-rank test). **K.** Average TTRs for Pre, MUS, and PBS sessions (mean ± SEM). Pre (13.42 ± 5.87) vs. MUS (148.08 ± 23.38), p = 0.002; MUS vs. Post (13.08 ± 5.70), p = 0.004; paired t-test. **L.** Scatter plot of TTRs vs. R2 correct rate across sessions. Each circle represents one session (open, Pre; red, MUS; black, PBS). Crosses denote group means ± SEM. **M-N.** Same as **I-J**, but for IC inactivation (N=7). Repeated measures two-way ANOVA; hit rate (drug, p = 6.21 x 10^-11^; rule, p = 0.005; interaction, p = 0.067); FA rate (drug, p = 0.566; rule, p = 0.409; interaction, p = 0.850); correct rate (drug, p = 2.99 x 10^-6^; rule, p = 0.044; interaction, p = 0.164). **O.** DiI-labeled brain sections showing multi-channel electrode tracks in the AC (top) and IC (bottom). AuD, dorsal secondary auditory cortex; Au1, primary auditory cortex; AuV, ventral secondary auditory cortex; DCIC, dorsal cortex of the inferior colliculus; CIC, central nucleus of the inferior colliculus; ECIC, external cortex of the inferior colliculus. Scale bar, 500 μm. **P.** Fraction of AC neurons (n=400) responsive to auditory stimuli during the stimulus epoch and task outcomes during the response window across R1, TR, and R2 phases. Black bars indicate neurons responsive across phases. Light blue (5kHz) and pink (10kHz) bars indicate stimulus-responsive neurons within each phase. Light gray bars indicate non-responsive neurons. Right, fraction of neurons responsive to 5 kHz only (light blue), 10 kHz only (pink), or both stimuli (light yellow) across task rules. Neurons were classified as responsive if they exhibited a significant stimulus-evoked response in at least one phase. Wilcoxon signed-rank test, p < 0.01. **Q.** Same as **P**, but for IC neurons (n=179). **R.** Left, example PSTHs from an AC neuron (#48) illustrating the gain modulation index (GMI) for the 5 kHz stimulus. Solid lines, R1; dotted line, R2; dark gray shading, stimulus epoch. Right, violin plots of population GMI values for 5 kHz-responsive AC neurons during TR and R2. Horizontal lines indicate medians across neurons responsive in at least one phase. Median GMI_5kHz_: TR = 0.055 (p = 0.044), R2 = 0.126 (p = 0.001); one-sample Wilcoxon signed-rank test with zeros. **S.** Same as **R,** but for IC neurons (#119). Median GMI_5kHz_: TR = -0.032, R2 = 0.004 (both n.s.). Between-group comparison: TR, p = 0.048.; R2, p = 0.016; Mood’s median test. **T-U**. Same as **R-S**, but for GMI_10kHz_. (**t**) AC median GMI_10kHz_: TR = 0.118 (p < 0.0001), R2 = 0.128 (p < 0.0001). (**u**) IC median GMI_10kHz_: TR = -0.022, R2 = -0.019 (both n.s.). Between-group comparison: TR, p = 0.001; R2, p = 0.0005. **V.** Schematic illustrating selectivity index (SI) calculation using ROC analysis. **W.** Example PSTHs from an AC neuron (#194) showing stimulus selectivity (5kHz versus 10kHz) during R1 (top) and R2 (bottom). SI values are indicated above each PSTH. **X.** Violin plots of changes in absolute stimulus selectivity (Δ |SI|) during R2 (|R2| - |R1|). Median Δ |SI|: AC = 0.056 (p = 0.026), IC = -0.003 (p = 0.542); Wilcoxon signed-rank test with zeros. Between-group comparison: p = 0.011; Mood’s median test. **Y.** Fraction of stable and flexible stimulus-selective neurons in AC and IC. Stable: AC = 0.180 ± 0.002, IC = 0.239 ± 0.002, p < 6.38 x 10^-8^; Flexible: AC = 0.261 ± 0.003, IC = 0.241 ± 0.004, p = 0.001; Wilcoxon rank-sum test.

On the first day of reversal training, mice exhibited high levels of false alarm (FA) rates in R2, continuing to lick in response to the previously rewarded but now incorrect stimulus (5 kHz), along with reduced hit rates for the newly rewarded stimulus (10 kHz; Figure 1B). With continued training, mice rapidly adjusted their licking behavior following rule reversal, achieving accurate performance within a single session (Figure 1C). Reversal-learning efficiency was quantified as the trial-number-to-reversal (TTR), defined as the number of trials required to reach expert-level performance in R2 (Methods). Across training sessions, R2 performance progressively improved, accompanied by a corresponding decrease in TTR, indicating enhanced reversal-learning efficiency over time (Figure 1D; Figure S1A). In expert mice, the average TTR was 30.82 ± 6.25 trials (Figure 1E). Lick reaction times were modestly increased in R2 relative to R1 (Figure 1F), whereas overall lick rates remained stable across rules (Figure 1G).

### Differential contributions of AC and IC during auditory reversal learning

To determine the causal contributions of auditory cortical and subcortical regions to rapid adaptation during auditory reversal learning, we inactivated the AC and IC using muscimol (MUS), a GABA_A_ receptor agonist, in expert mice performing the task (Figure 1H; Figures S1B and Table S1; Methods). Bilateral AC inactivation selectively impaired auditory reversal learning, as evidenced by increased TTRs, reduced Hit rates, and elevated FA rates in R2, while leaving performance in R1 intact (Figures 1I-1L; Figures S1C, S1F and S1G). In contrast, unilateral IC inactivation disrupted auditory discrimination performance in both R1 and R2, significantly reducing hit rates across task phases, including R1 (Figures 1M and 1N; Figures S1D, S1H-S1I). Control PBS injections into either AC or IC had no effect on behavior performance (Figures 1I-1N; Figures S1C, S1D, S1F-S1I).

Given these distinct behavioral effects, we next examined how auditory representations in AC and IC evolve across task rules during reversal learning. We recorded single-unit activity from AC (n = 400 neurons) and IC (n = 179 neurons) using linear silicon probes in mice performing the auditory reversal-learning task (three mice per region, with two recorded in both regions; Figure 1O; Table S2). Behavioral performance, including correct rates, TTRs, reaction times, and lick rates, did not differ between recording groups (Figures S2A-S2E; Supplementary Table 2). Auditory-evoked activity was analyzed across three task phases: R1 (initial rule), TR (transition from R1 to R2 up to TTR), and R2 (reversed rule after TTR). To minimize motor-related confounds, analyses were restricted to trials without licking within ±1 s of stimulus onset. Neurons exhibiting significant responses to either the 5 kHz or 10 kHz tone in each phase were identified using a Wilcoxon signed-rank test (p < 0.01). The fraction of stimulus-responsive neurons decreased during TR but was comparable between R1 and R2 in both AC and IC (Figures 1P and 1Q).

We first examined whether stimulus-responsive neurons altered their responses to the same auditory stimuli across task phases (Figures 1R-1U). Changes in response amplitude were quantified using the auditory gain modulation index (GMI), where positive values indicate enhanced responses and negative values indicate suppressed responses relative to R1. AC neurons exhibited significantly enhanced auditory responses in R2 relative to R1 for both tones (Figures 1R and 1T), indicating an overall amplification of auditory representations following rule reversal. Notably, these enhanced responses emerged during the transition phase, suggesting that gain modulation develops early during behavioral adaptation in the AC (Figures 1R and 1T). In contrast, IC neurons exhibited stable tone-evoked responses across task phases (Figures 1S and 1U).

We next examined whether tone selectivity changed across task phases. The tone selectivity index (SI) was quantified for each neuron using receiver operating characteristic (ROC) analysis^23,31^, based on differences in firing rates during the stimulus epoch between the 5kHz and 10kHz tones (Figures 1V and 1W; Methods). Stimulus selectivity was significantly enhanced in R2 exclusively in AC neurons, with no corresponding change in IC neurons (Figure 1X). Consistently, a larger fraction of AC neurons exhibited changes in their auditory representations across rules compared to IC neurons (Figure 1Y, Figures S2F and S2G). Together, these results indicate that AC neurons undergo pronounced remapping of auditory representations following rule reversal, whereas IC neurons maintain stable auditory encoding across task rules.

Collectively, these findings reveal a functional dissociation between auditory cortical and subcortical regions during auditory reversal learning. The AC is specifically required for behavioral flexibility following contingency reversal and exhibits enhanced representation of reversed Go/No-go stimuli through integration of rule information with auditory inputs. In contrast, the IC supports stable auditory discrimination performance by maintaining reliable auditory encoding and consistent execution of learned responses across task phases.

### PPC, projecting to AC and IC, supports auditory reversal learning

To identify candidate top-down brain regions that balance flexibility in the AC and stability in the IC during auditory reversal learning, we performed dual-color retrograde tracing by injecting retrograde adeno-associated viruses (AAVrg) expressing GFP and tdTomato into the AC and IC, respectively (Figure 2A). Whole-brain clearing combined with light-sheet microscopy revealed distinct anatomical projection patterns to each structure (Figures 2A-2C; Supplementary Movies 1 and 2). The orbitofrontal cortex (OFC), secondary motor cortex (M2), and cingulate cortex area 1 (Cg1) preferentially projected to the AC (Figure 2C_i__-ii_), whereas the medial prefrontal cortex, including prelimbic (PrL) and infralimbic (IL) cortices, as well as cingulate cortex area 2 (Cg2), primarily innervated the IC (Figure 2c_i__-ii_; Supplementary Movie 2). The insular cortex (InC), extending along the anterior-posterior axis, mainly projected to the IC, whereas the adjacent claustrum (Cl) exhibited strong projections to the AC (Figure 2C_ii__-iii_).

**Figure 2.**
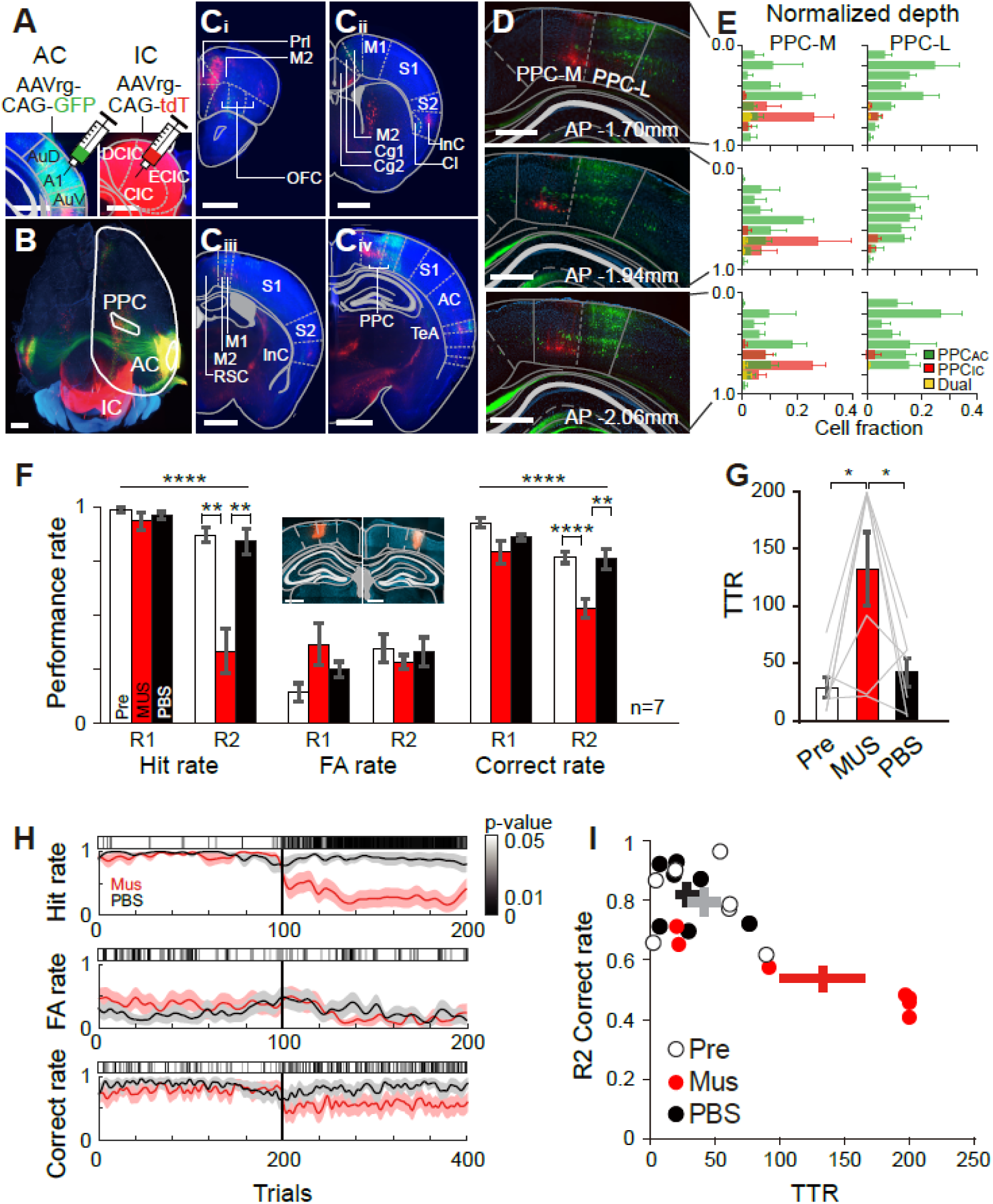
PPC sends segregated top-down projections to AC and IC and is required for auditory reversal learning. **A-E.** Dual-color retrograde tracing reveals anatomically segregated PPC projections to the AC (green) and IC (red). **A.** Coronal view of a cleared whole brain showing retrograde injection sites in the AC (AAVrg-CAG-GFP, left) and IC (AAVrg-CAG-tdTomato, right), aligned to the Paxinos atlas (gray lines). Scale bars, 500 μm. **B.** Dorsal view of a cleared brain imaged with light-sheet microscopy and reconstructed in 3D. White outlines indicate the right cortical hemisphere and PPC. GFP-labeled PPC_AC_ neurons are positioned laterally, whereas tdTomato-labeled PPC_IC_ neurons are located medially. Scale bar, 1 mm. **C.** Coronal sections from cleared brains imaged by light-sheet microscopy and reconstructed in 3D. DAPI (blue) indicates brain architecture; AC-projecting (green) and IC-projecting (red) neurons are visualized across cortical regions. Images aligned to the Paxinos atlas (gray lines). Scale bars, 1mm. **C_i_**, OFC (orbitofrontal cortex; medial, ventral, and lateral parts); Prl (prelimbic cortex); M2 (secondary motor cortex). **C_ii_**, M1 (primary motor cortex); M2; S1 (primary somatosensory cortex); S2 (secondary somatosensory cortex); Cg1 (cingulate cortex area 1); Cg2 (cingulate cortex area 2); InC (insular cortex); Cl (claustrum). **C_iii_**, M1; M2; S1; S2; InC. **C_iv_,** PPC (medial and lateral); S1; AC; TeA (temporal association cortex). D. Representative coronal sections along the anterior-posterior (AP) axis of the PPC showing segregated PPC_AC_ (green) and PPC_IC_ (red) neuronal populations. Images are aligned to the Paxinos atlas (gray lines). Scale bars, 500 μm. **E.** Laminar and spatial distribution of retrogradely labeled neurons in medial PPC (PPC-M, left) and lateral PPC (PPC-L, right). Neuronal fractions were binned by normalized cortical depth (pia = 0, ventricle = 1) using DAPI signals. Green, PPC_AC_ neurons; red, PPC_IC_ neurons; yellow, dual-labeled neurons. **F-I.** PPC inactivation impairs auditory reversal learning. F. Behavioral effects of PPC inactivation (N=7). Average hit, FA, and correct rates during Pre (no injection; open bars), MUS (red), and PBS (black) sessions. Repeated-measures two-way ANOVA; hit rate: drug, p = 3.84 x 10^-9^; rule, p = 1.83 x 10^-7^; interaction, p = 3.45 x 10^-8^; FA rate: drug, p = 0.666; rule, p = 0.423; interaction, p = 0.050; Correct rate: drug, p = 0.0001; rule, p = 0.001; interaction, p = 0.016. Inset, bilateral PPC injection sites labeled with fluoro-ruby and aligned to the Paxinos atlas. Scale bars, 500 μm. G. Average TTRs for Pre (28.92 ± 9.08), MUS (133.21 ± 32.37), and PBS (42.29 ± 12.55) sessions (mean ± SEM). Paired t-test: Pre vs. MUS, p = 0.019; MUS vs. Post; p = 0.034. H. Session-averaged performance with or without PPC inactivation (N=7). hit (top); FA (middle), correct (bottom) rates are shown for MUS (red) and PBS (black) sessions. Shaded regions indicate ± SEM. Gray-scaled bars indicate p-values for significant differences between MUS and PBS sessions; Wilcoxon signed-rank tests. I. Scatter plot of TTRs vs. R2 correct rate across individual sessions. Circles, individual sessions (open, Pre; red, MUS; black, PBS). Crosses denote group means ± SEM.

In contrast to these regions, which selectively projected to either the AC or IC, both the PPC and temporal association cortex (TeA), corresponding to dorsal and ventral association cortices, respectively, contained two anatomically distinct and spatially segregated neuronal populations projecting to the AC and IC (Figures 2B and 2C_iv_; Supplementary Movies 1 and 2). In particular, robust dual labeling in the PPC indicated prominent top-down projections to both auditory areas (Figures 2B, 2D and 2E). Within the PPC, neurons projecting to the IC (PPC_IC_) accounted for 19.30 % ± 4.69% of labeled cells, whereas neurons projecting to the AC (PPC_AC_) comprised 82.03 % ± 4.61% (Figure S3). Neurons co-labeled with both tracers were rare, constituting only 1.66 % ± 0.31% of PPC_AC_ neurons and 7.85% ± 1.93% of PPC_IC_ neurons. Spatially, PPC_AC_ neurons were distributed along the mediolateral axis with higher density in lateral PPC (PPC-L), whereas PPC_IC_ neurons were predominantly localized to medial PPC (PPC-M) (Figures 2B, 2C_iv,_ and 2D). Depth analysis using 4′,6-diamidino-2-phenylindole (DAPI)-stained sections revealed that PPC_AC_ neurons spanned superficial to deep cortical layers, whereas PPC_IC_ neurons were largely confined to deep layers (Figures 2D and 2E; Methods). These anatomical distinctions are consistent with previously described intratelencephalic (IT) and pyramidal tract (PT) cortical neuron subtypes^50,51^.

We next examined the functional contribution of the PPC to auditory reversal learning. Similar to the effects of AC inactivation (Figures 1I-1L; Figures S1C, S1F, and S1G), bilateral PPC inactivation using muscimol significantly increased TTRs following the rule switch and reduced hit rates in R2 (Figures 2F-2I; Figures S1E, S1J and S1K). Importantly, this deficit was specific to reversal learning, as auditory discrimination performance during R1 remained intact (Figures 2F and 2H; Figures S1E, S1J and S1K). Control PBS injections had no effect on TTRs or task performance (Figures 2F-2I; Figures S1E, S1J and S1K). Together, these results demonstrate that PPC activity is required for flexible adaptation to reversed stimulus-outcome contingencies during auditory reversal learning.

### Double dissociation of top-down projections from the PPC during reversal learning

Given that segregated PPC neuronal populations project to the AC and IC, we next examined the functional roles of these two top-down pathways in auditory reversal learning. To selectively inhibit each projection during task performance, we expressed the inhibitory opsin archaerhodopsin (ArchT) in PPC_AC_ or PPC_IC_ neurons via bilateral injections of AAVrg expressing ArchT under the CaMKIIa promoter into the AC or IC, respectively (Figures 3A and 3B; Table S3). Green laser illumination was delivered to the PPC from 1 s before stimulus onset through the response window either selectively during the transition phase (TR; early R2 up to the TTR; Type I inactivation) or on randomly selected trials during stable phases (R1 or R2; Type II inactivation).

**Figure 3.**
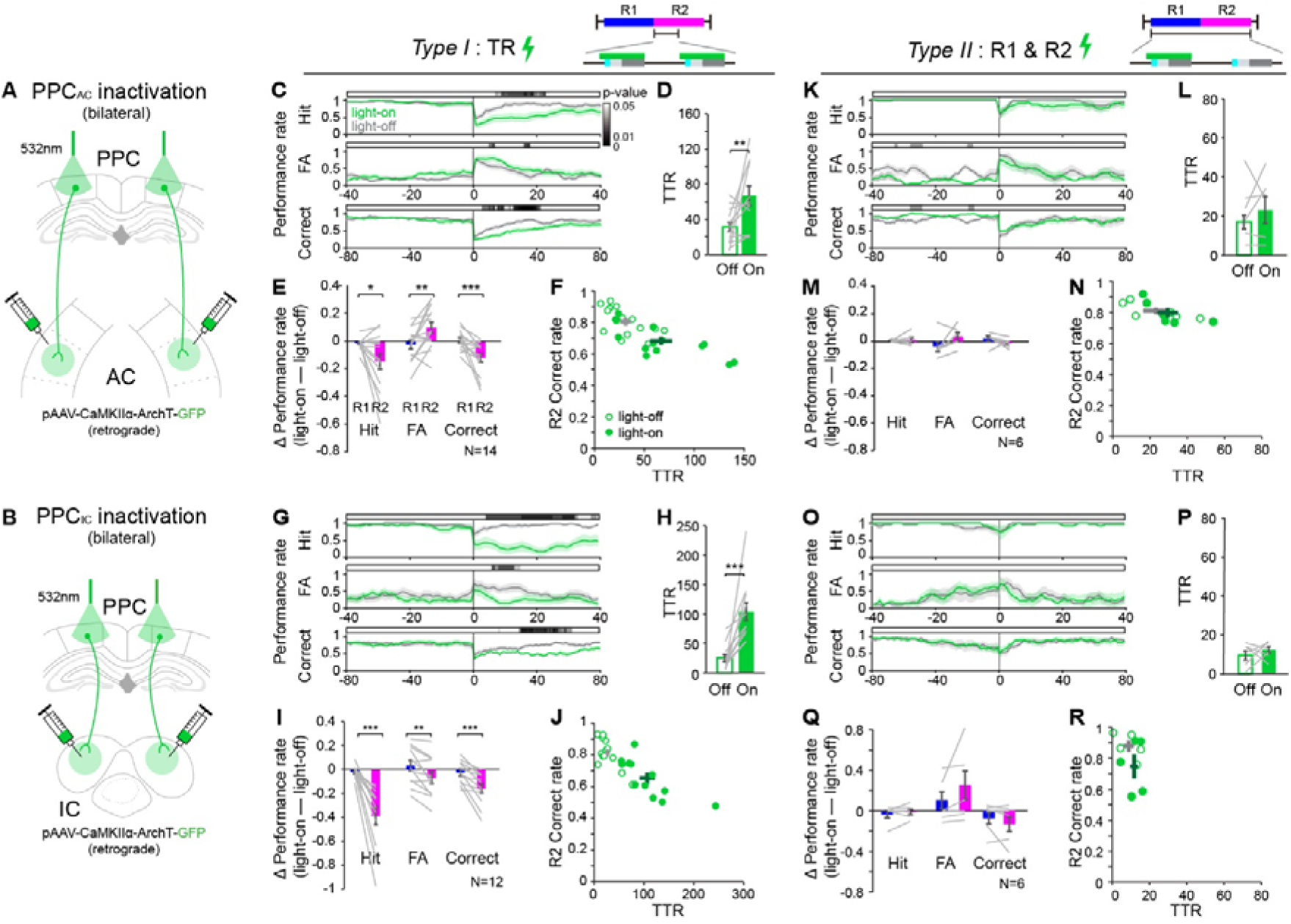
Double dissociation of PPC top-down projections during auditory reversal learning. **A-B.** Schematics of projection-specific optogenetic inactivation targeting PPC_AC_ neurons (**A)** and PPC_IC_ neurons (**B**) **C-F**. Impaired reversal learning induced by Type I PPC_AC_ inactivation (N=14 mice). Top, Type I inactivation protocol. Green laser light (532 nm) was delivered immediately after rule reversal, from 1 s before stimulus onset to the end of the response window. **C.** Time course of hit (top), FA (middle), and correct (bottom) rates aligned to the start of R2 (trial 0). Green, light-on sessions; gray, light-off sessions; shading, ± SEM. Gray-scaled bars indicate p-values comparing light-off and light-on sessions; Wilcoxon signed-rank test. **D.** TTRs were significantly increased during light-on (66.79 ± 10.98) compared to light-off (31.29 ± 5.37) sessions; p = 0.009, Wilcoxon signed-rank test. **E.** Changes in performance (Δ = light-on - light-off) for hit (R1, -0.015 ± 0.011; R2, -0.148 ± 0.057; p = 0.042), FA (R1 = -0.027 ± 0.032; R2 = 0.096 ± 0.040; p = 0.007), and correct rates (R1, 0.005 ± 0.019; R2, -0.167 ± 0.052; p = 0.001). Wilcoxon signed-rank test. **F.** Scatter plot of TTR vs. R2 correct rates across sessions. Filled green circles, light-on; open green circles, light-off. Crosses show group means ± SEM (dark-green, light-on; gray, light-off). **G-J.** Same as **C-D**, but for Type I PPC_IC_ inactivation (N=12). **H.** TTR was significantly increased in light-on (104.08 ± 14.95) compared to light-off sessions (25.08 ± 5.94); p = 0.0005, Wilcoxon signed-rank test. **I.** Change in performance for hit (R1, -0.022 ± 0.015; R2, -0.388 ± 0.077; p = 0.0005), FA (R1 = 0.032 ± 0.045; R2 = -0.069 ± 0.049; p = 0.003), and correct rates (R1, -0.027 ± 0.027; R2, -0.160 ± 0.038; p = 0.0005). Wilcoxon signed-rank test. **K-N.** Same as **C**-**F** but for Type II PPC_AC_ inactivation (N=6). Top, Type II inactivation protocol. Laser light was delivered randomly in 50% of trials across all task phases, from 1 s before stimulus onset to the end of the response window. Note that mice showed intact reversal learning. **L.** TTRs did not differ between light-on (29.33 ± 5.58) and light-off (23.00 ± 6.84) sessions. **M.** Change in performance rates for hit (R1, 0.003 ± 0.003; R2, 0.010 ± 0.018; n.s.), FA (R1 = -0.036 ± 0.037; R2 = 0.0.35 ± 0.030; n.s.), and Correct rates (R1, 0.019 ± 0.019; R2, -0.012 ± 0.014; n.s.). **O-R.** Same as **C**-**F,** but for Type II PPC_IC_ inactivation (N=6). **P.** TTR did not differ between light-on (9.40 ± 2.29) and light-off (12.40 ± 1.74) sessions. **Q.** Change in performance for hit (R1, -0.035 ± 0.035; R2, -0.010 ± 0.026; n.s.), FA (R1 = 0.105 ± 0.085; R2 = 0.253 ± 0.138; n.s.), and correct rates (R1, -0.072 ± 0.057; R2, -0.131 ± 0.074; n.s.).

Type I inactivation of PPC_AC_ neurons induced pronounced perseverative errors, reflected by decreased Hit rates, increased FA rates, and elevated TTR values (Figures 3C-3F), indicating a failure to update reversed stimulus-outcome contingencies. In contrast, Type I inactivation of PPC_IC_ neurons markedly reduced licking following rule reversal and resulted in a significant decrease in Hit rates, indicating an inability to maintain Go responses after the contingency switch (Figures 3G-3J). Control mice expressing eYFP in the PPC showed no behavioral effects under the same stimulation protocol (Figure S4), confirming the specificity of these effects. By comparison, Type II inactivation of either pathway had no detectable impact on task performance across rules (Figures 3K-3R), indicating that neither PPC_AC_ nor PPC_IC_ activity is required once mice have fully adapted to stable contingencies. Together, these results indicate that both PPC output pathways are specifically engaged during contingency reversal, with PPC_AC_ supporting contingency updating and PPC_IC_ supporting stable action execution.

Collectively, projection-specific optogenetic perturbations reveal a double dissociation between PPC_AC_ and PPC_IC_ function during reversal learning. Inactivation of PPC_AC_ disrupts contingency updating, leading to perseverative licking responses to previously rewarded stimuli that have become No-go. In contrast, inactivation of PPC_IC_ impairs the execution of Go responses, reflecting a failure to sustain goal-directed licking behavior following the rule change.

### PPC_AC_ exhibits flexible reorganization of population activity, whereas PPC_IC_ remains stable during reversal learning

We next investigated how the two PPC output pathways differentially support adaptive behavior during reversal learning at the population level. PPC neurons projecting to either the AC or IC were selectively labeled by injecting retrograde AAVs expressing Cre recombinase into each target region in Ai148 mice expressing GCaMP6f (Figures 4A and 4B; Methods). This approach yielded 283 PPC_AC_ neurons and 111 PPC_IC_ neurons with sufficient activity for analysis. Behavioral performance, including correct rates, TTRs, lick rates, and reaction times across rules and trial types, did not differ between the two imaging groups (Figures S5A-S5E; Table S4).

**Figure 4.**
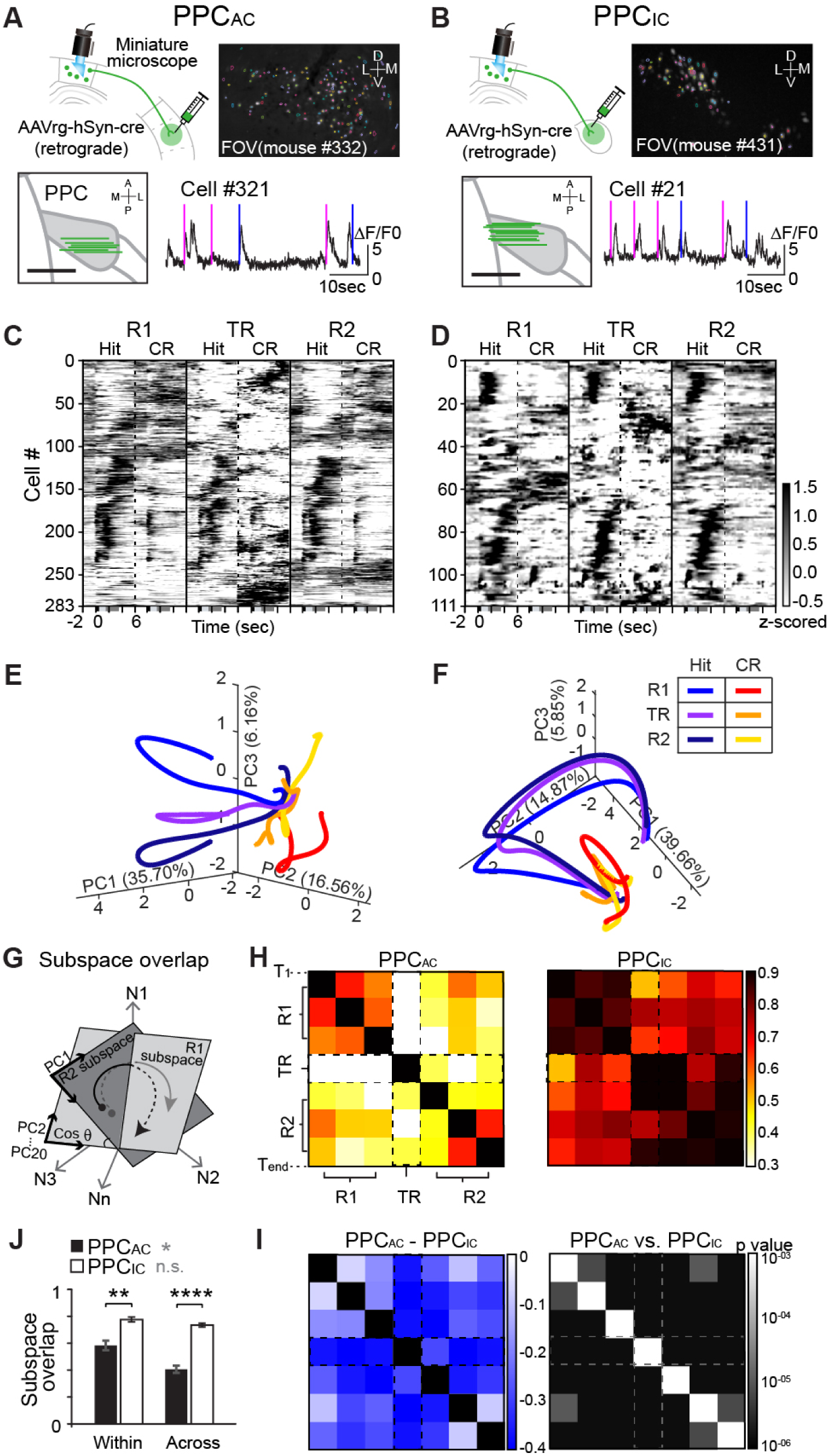
Distinct population dynamics of PPC_AC_ and PPC_IC_ neurons during auditory reversal learning. **A.** Projection-specific Ca^2+^ imaging of PPC_AC_ neurons. Top left, schematic of the imaging strategy. Top right, representative field of view (FOV) in the PPC. Bottom left, dorsal view showing all FOV locations. Bottom right, example Ca^2+^ trace from a single neuron during task performance. Vertical blue and magenta lines indicate 5kHz and 10kHz stimulus onset, respectively. Scale bar, 1mm. **B.** Same as **A**, but for PPC_IC_ neurons. **C.** Rastermap visualization (50 clusters) of trial-averaged, z-scored Ca^2+^ activity from PPC_AC_ neurons (n = 283) during concatenated hit and CR trials across R1 (left), TR (middle), and R2 (right). Neurons were clustered based on activity during R1. **D.** Same as **C**, but for PPC_IC_ neurons (n=111). **E-F.** Principal component trajectories of trial-averaged population activity from PPC_AC_ (**E**) and PPC_IC_ (**F**) neurons, aligned to stimulus onset (-2 to +6 s) and projected onto the first three PCs. Colors indicate trial type and task phase, as shown in the legend. **G.** Schematic of the subspace-overlap analysis, computed from angular differences between PC axes. **H.** Subspace-overlap heatmaps for PPC_AC_ (left) and PPC_IC_ (right) populations across task phases. Color scale: 0 (no overlap) to 1 (complete overlap). **I.** Left, difference in subspace overlap (PPC_AC_ - PPC_IC_), showing greater overlap in PPC_IC_ neurons, particularly during TR. Right, statistical comparison of overlap values between PPC_AC_ and PPC_IC_ populations across 20-fold cross-validation (Wilcoxon rank-sum test). **J.** Average subspace overlaps (mean ± SEM) within and across task phases for PPC_AC_ (black bars; within, 0.601 ± 0.036; across, 0.419 ± 0.028) and PPC_IC_ neurons (open bars; within, 0.799 ± 0.019; across, 0.756 ± 0.014). PPC_AC_ neurons exhibited a significantly greater reduction in overlap across phases compared to within phases (p = 0.0072), whereas PPC_IC_ neurons showed no within-group difference (Wilcoxon rank-sum test). Between-group comparisons revealed significantly lower overlaps in PPC_AC_ than PPC_IC_ populations both within (p = 0.0022) and across phases (p = 3.39 x 10^-6^; Wilcoxon rank-sum test).

To visualize population activity across task phases, we applied Rastermap, a scaled k-means clustering algorithm^52,53^, to the mean activity of neurons during Hit and CR trials and plotted cluster-ordered activity across R1, TR, and post-transition R2 (Figures 4C and 4D). PPC_AC_ neurons exhibited more robust activity reorganization compared with PPC_IC_ neurons. To further characterize population dynamics, we performed principal component analysis (PCA; Methods) and projected activity trajectories for hit and CR trials from R1, TR, and R2 into a shared low-dimensional space (Figures 4E and 4F; Figure S5F). PPC_AC_ population trajectories showed clear remapping across phases, with divergence reflecting both contingency-dependent trial types (hit vs. CR) and task rules (R1 vs. R2; Figure 4E). In contrast, PPC_IC_ population activity was primarily driven by hit trials and remained largely stable, with minimal trajectory shifts following rule reversal (Figure 4F).

We next quantified population-level changes using subspace overlap analysis (Figure 4G; Methods). Both PPC_AC_ and PPC_IC_ populations exhibited strong within-rule overlap, indicating consistent activity under stable contingencies. However, PPC_AC_ neurons showed significantly reduced overlap across task phases (Figure 4H), consistent with dynamic reorganization during reversal learning. By contrast, PPC_IC_ neurons maintained high subspace overlap across all phases (Figure 4H), and across-rule overlap was significantly greater in PPC_IC_ than in PPC_AC_ populations (Figures 4I and 4J). Together, these results demonstrate a functional dissociation between PPC output pathways: PPC_AC_ neurons flexibly reorganize their population activity to encode updated stimulus-outcome relationships, whereas PPC_IC_ neurons preserve stable activity patterns during Go trials, potentially supporting reliable execution of learned actions across rule reversals.

### PPC top-down projections encode distinct task variables during the reversal transition

To determine how PPC_AC_ and PPC_IC_ populations encode task variables (TVs) during the reversal transition (TR), when stimulus-outcome updating occurs, we fit a generalized linear model (GLM; Methods) to single-neuron activity across phases. TVs included stimulus identity (5kHz, 10kHz), licking behavior (bout onset and offset), and trial outcomes (hit, FA, CR, miss; Figures 5A and 5B). Approximately half of PPC_AC_ and PPC_IC_ neurons encoded at least one TV across phases, with PPC_IC_ neurons exhibiting overall stronger model fits (Figure 5A; Figures S6A and S6B).

**Figure 5.**
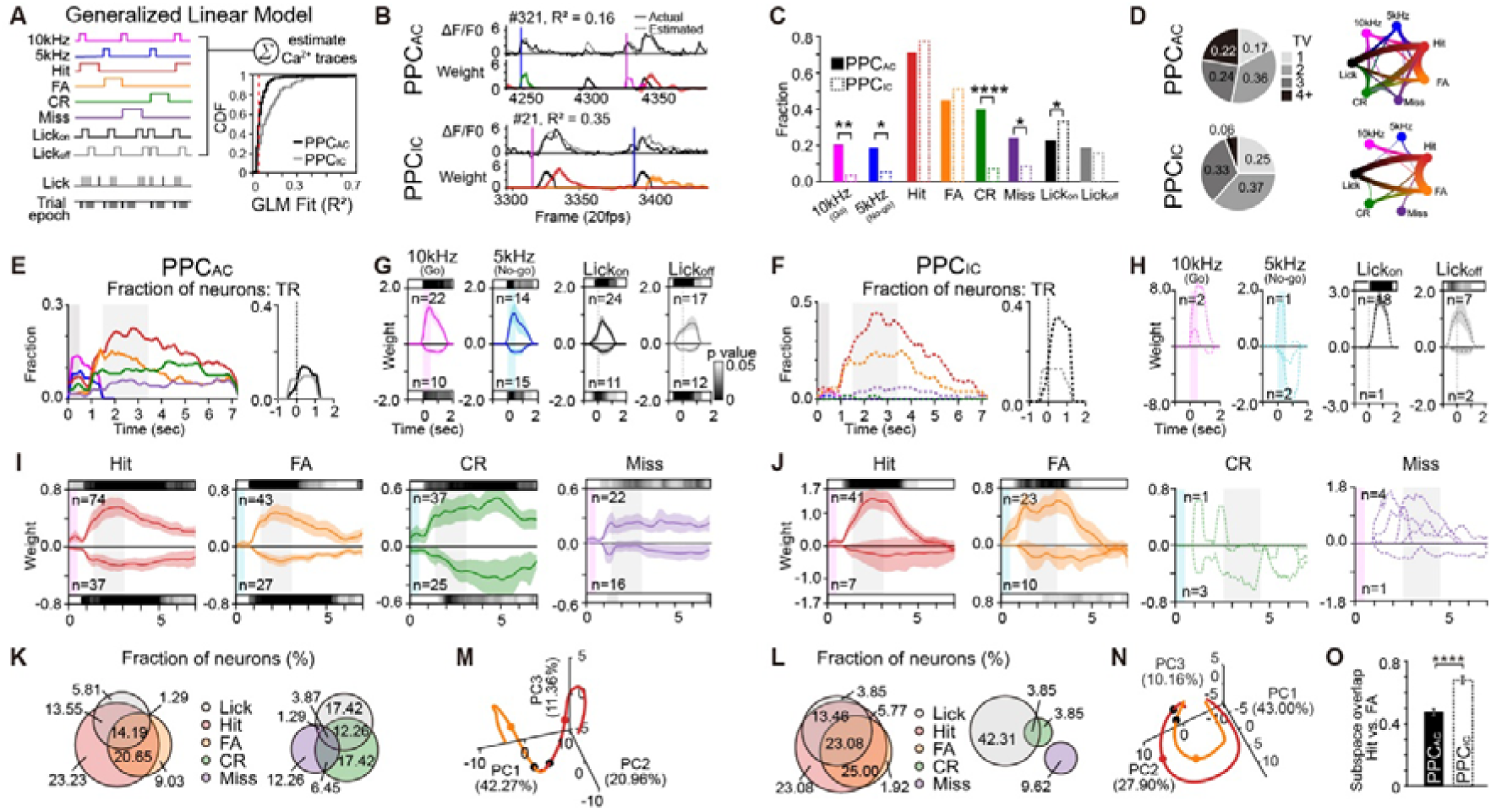
Distinct and multiplexed task-variable encoding by PPC_AC_ and PPC_IC_ neurons during reversal learning. **A.** Left, schematic of the GLM used to quantify encoding of task variables (TVs) by individual neurons. Right, cumulative distribution of GLM-fitted R^2^ values during the TR for PPC_AC_ (black) and PPC_IC_ (gray) neurons. **B.** Example Ca^2+^ traces with corresponding GLM fits and TV weights from representative PPC_AC_ (top) and PPC_IC_ (bottom) neurons. Solid lines indicate measured Ca^2+^ activity; dotted lines indicate GLM-predicted activity. Vertical blue and magenta lines indicate onset of the 5kHz and 10kHz stimuli, respectively. TV weights are color-coded as in **A**. **C.** Fraction of neurons encoding each TV during TR. Filled bars, PPC_AC_; open bars, PPC_IC_. Two-sided chi-square tests; 10kHz, p = 0.004; 5kHz, p = 0.026; CR, p < 0.0001; miss, p = 0.022; Lick_on_, p = 0.044; hit, FA and Lick_off_, not significant. **D.** Left, fraction of PPC_AC_ (top) and PPC_IC_ (bottom) neurons grouped by the number of TVs encoded. Right, circular Sankey diagrams illustrating multiplexed TV co-encoding in PPC_AC_ (top) and PPC_IC_ (bottom) populations. Line thickness indicates frequency of each combination. **E-F.** Time-resolved fractions of neurons encoding each TV across the trial for PPC_AC_ (**E**) and PPC_IC_ (**F**) neurons. Left, stimulus- and outcome-encoding from 0 to 7 s relative to stimulus onset. Dark and light gray shading denote stimulus and response epochs, respectively. Right, fractions of neurons encoding motor variables (lick-bout onset and offset, aligned to 0 s). Task variables are color-coded as in **a**. **G.** Average GLM weights for PPC_AC_ neurons encoding 10kHz (Go, magenta), 5kHz (No-go, blue), Lick_on_ (black), and Lick_off_ (gray) variables. Positive (top) and negative (bottom) weights are shown separately; shading indicates ± SEM. Gray-scaled bars indicate significance (one-sample Wilcoxon signed-rank test). **H**. Same as **G**, but for PPC_IC_ neurons. When fewer than five neurons encoded a given variable, individual traces are plotted as dotted lines. **I-J**. Average GLM weights for neurons encoding outcome variables (hit, FA, CR, and miss) in PPC_AC_ (**I**) and PPC_IC_ (**J**). **K-L**. Proportional Euler diagrams showing overlap among PPC_AC_ (**K**) and PPC_IC_ (**L**) neurons encoding hit, FA, and lick variables (left), and CR, miss, and lick variables (right). **M-N**. PCA trajectories of hit - and FA-encoding PPC_AC_ (**M**) and PPC_IC_ (**N**) neurons during hit and FA trials. Black dots denote stimulus onset; colored dots indicate response-window onset. **O**. Quantification of trajectory separation using subspace-overlap analysis. Wilcoxon signed-rank test, p = 6.796 x 10^-8^.

Despite comparable encoding prevalence, the structure of TV representation differed between pathways. PPC_AC_ neurons encoded a greater number of TVs per neuron and exhibited more multiplexed co-encoding across variables, whereas PPC_IC_ neurons were more selectively tuned to licking-related variables and lick-associated outcomes (Figures 5C and 5D). PPC_AC_ neurons showed stronger encoding of auditory stimulus identity and broadly represented trial outcomes, including both lick-associated (hit and FA) and no-lick-associated (CR and miss), whereas PPC_IC_ neurons preferentially encoded licking behavior and lick-associated outcomes (hit and FA; Figures 5C-5H). Both populations increased encoding of task-relevant outcomes during TR relative to R1 and R2, consistent with engagement during contingency updating (Figures 5E and 5F; Figures S6D and S6E). However, PPC_AC_ neurons showed greater instability of task-variable encoding across phases, with fewer neurons maintaining stable representations through TR compared to PPC_IC_ neurons (Figure S6C).

Temporal analysis of GLM weights revealed further dissociation. PPC_AC_ neurons exhibited prolonged and distributed encoding of both lick and no-lick outcomes across the trial, whereas PPC_IC_ neurons encoded lick-related outcomes predominantly within the response window (Figures 5I and 5J). Hit and FA encoding largely overlapped with lick encoding in both populations but more strongly in PPC_IC_ neurons, while encoding of no-lick outcomes showed minimal overlap with lick-related variables (Figures 5K and 5L). Notably, PPC_AC_ neurons differentiated rewarded and unrewarded lick outcomes, with divergent activity patterns for Hit versus FA trials, whereas PPC_IC_ neurons showed similar activity across these conditions (Figures 5M and 5O). Sensory encoding in PPC_AC_ was largely segregated from motor and outcome representations, indicating parallel processing of auditory information within this pathway (Figure S6F).

Together, these results demonstrate that PPC_AC_ neurons flexibly and dynamically encode newly relevant stimulus-outcome relationships during reversal, while PPC_IC_ neurons maintain stable representations of licking and lick-associated outcomes, supporting consistent action execution across changing contingencies.

### A feedforward network model with parallel PPC pathways for rapid reversal learning

To examine the circuit mechanisms underlying the flexibility-stability trade-off during reversal learning, we developed a hierarchical feedforward network model linking sensory input to action decision. The network architecture reflects the biological hierarchy of auditory processing, with sensory signals transmitted from the IC to the AC, and convergent projections from both regions to a downstream motor unit that generates lick or no-lick decisions (Figures 6A and 6B). In the model, the IC provides a stable auditory-to-action drive, whereas the AC implements rule-dependent plasticity to update lick/no-lick associations.

**Figure 6.**
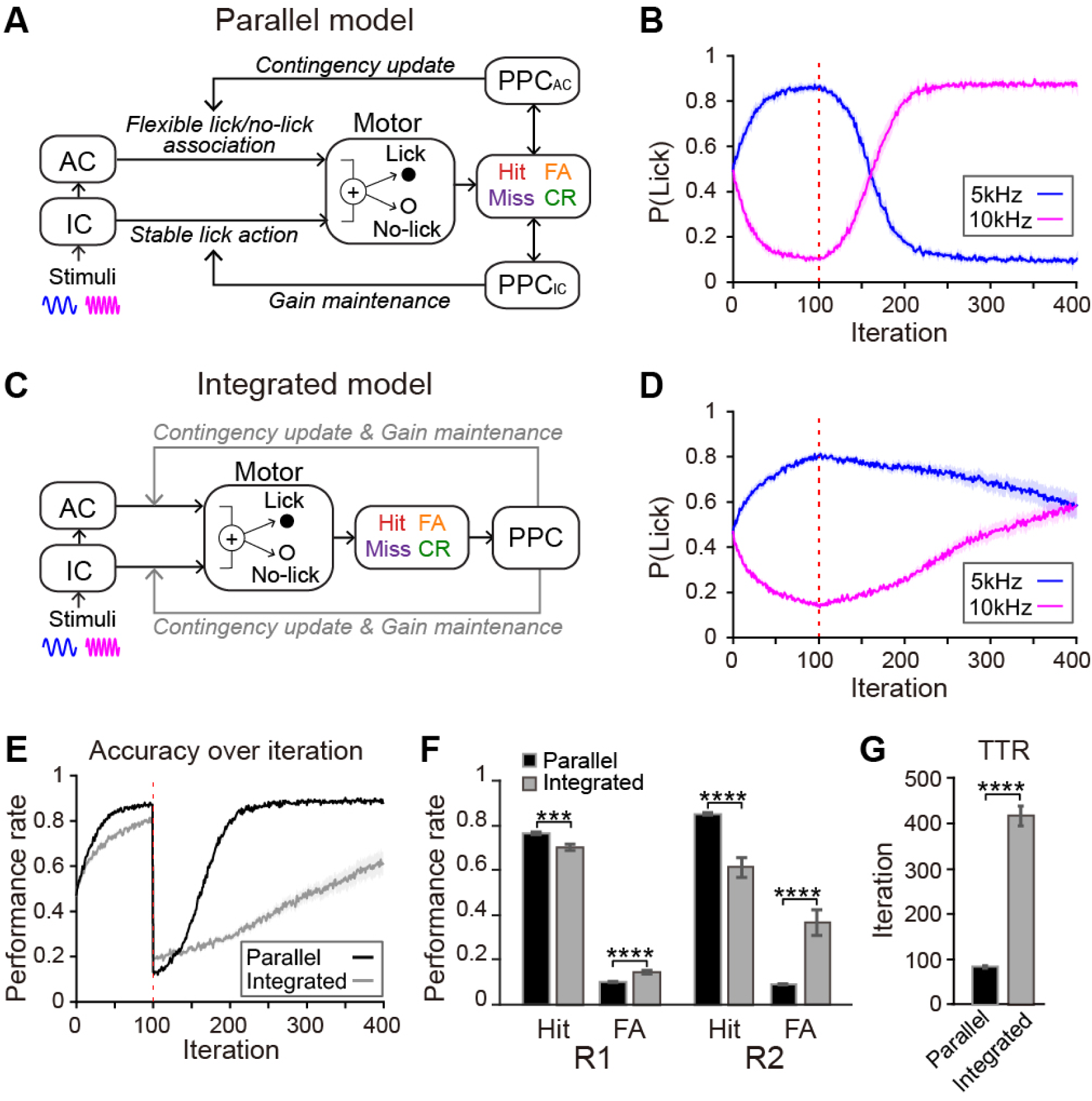
Computational dissociation of flexibility and stability *via* parallel PPC circuits. **A.** Schematic of the parallel PPC top-down model. Auditory input is processed by the IC layer and relayed to the AC layer. Activity in both layers converges to drive motor output (lick or no-lick). In this architecture, the PPC_AC_ module regulates contingency-dependent plasticity in AC, whereas the PPC_IC_ module stabilizes auditory-to-lick output in the IC through goal-directed signals. **B.** Evolution of lick probabilities for the two auditory stimuli across learning in the parallel top-down circuit. The vertical dashed red line indicates the rule-reversal point. **C.** Schematic of an integrated PPC model, in which a single PPC module jointly monitors contingency and goal errors to regulate both AC plasticity and IC gain. **D.** Same as **B**, but for the integrated top-down circuit. **E.** Learning curves showing task accuracy for the parallel (black) and integrated (gray) networks. The parallel network exhibits faster recovery following reversal, indicating more rapid adaptation. **F.** Comparison of behavioral performance between parallel and integrated networks during R1 (iterations 1-100) and R2 (iterations 300-400). Wilcoxon rank-sum test; R1 hit rate (p = 0.0003); R1 FA rate (p = 0.0003); R2 hit rate (p < 0.00001); R2 FA rate (p < 0.00001). **G.** TTR was significantly higher in the integrated network (83.300 ± 2.264) than in the parallel network (416.700 ± 22.263); p < 0.00001, Wilcoxon rank-sum test.

These two pathways are independently modulated by parallel PPC control modules, PPC_IC_ and PPC_AC_, whose signals evolve dynamically across task phases. During initial learning (R1), PPC_AC_ contingency error signals decrease as stimulus-outcome associations are acquired, while PPC_IC_ output stabilizes to support consistent action execution. Following rule reversal (R2), PPC_AC_ detects a sharp increase in contingency error and transiently gates elevated plasticity in the AC, enabling rapid overwriting of outdated associations. In parallel, PPC_IC_ maintains stable feedback to the IC pathway, preserving effective sensory-to-motor transformation and sustaining task engagement despite transiently unreliable outcomes.

The functional roles of each pathway were validated through simulated circuit-specific inactivation (Figure S7). Disabling the PPC_AC_ pathway selectively impaired contingency updating, reducing behavioral flexibility, whereas disabling the PPC_IC_ pathway disrupted action stability, reducing behavioral consistency. To test whether pathway segregation is required, we simulated an integrated architecture in which a single control module jointly regulated both plasticity and action stability (Figures 6C and 6D). The integrated model failed to balance flexibility and stability, resulting in slower reversal learning and reduced synaptic plasticity following contingency reversal (Figures 6B, 6D, and 6E-6G). Together, these simulations demonstrate that segregation of PPC top-down control into parallel pathways is necessary to resolve the fundamental dilemma between behavioral persistence and flexibility, enabling rapid and stable adaptation following contingency reversal.

## Discussion

Behavioral flexibility requires the ability to update internal models in response to changing rules while maintaining stable, goal-directed actions. Although prefrontal and parietal regions have long been implicated in reversal learning and executive control^2,54,55^, the circuit mechanisms that balance flexibility and stability remain poorly defined. Here, we identify a circuit-level organizational principle in the mouse brain in which anatomically segregated top-down projections from the PPC to AC and IC play complementary roles during auditory reversal learning (Figure 6A). PPC projections to AC (PPC_AC_) flexibly reorganize their activity to encode stimulus-outcome contingencies and support rule updating, whereas projections to IC (PPC_IC_) maintain stable representations related to goal-directed licking execution. This double dissociation reveals how parallel PPC circuits coordinate flexible updating with stable action control.

This circuit architecture suggests a general top-down motif for resolving the flexibility-stability trade-off through parallel control. By anatomically segregating these outputs, the PPC minimizes interference between adaptive and stable computations. In a unified network, the rapid remapping of task rules could inadvertently disrupt the signals required for stable motor execution, leading to premature stopping or a collapse of behavioral drive, a phenomenon we observed during PPC_IC_ inactivation. Thus, parallel gating serves as a hardware solution to the fundamental challenge in cognitive systems: integrating new information without overwriting previously acquired execution policies^56^. This buffering mechanism allows biological systems to maintain high behavioral drive (via PPC_IC_) even when the internal model of the environment is in a state of high uncertainty (during PPC_AC_-mediated remapping).

Our analysis of PPC_AC_ neural dynamics further reveals a coding strategy that combines model-based and model-free reinforcement learning. On one hand, PPC_AC_ populations exhibit systematic shifts in their activity patterns that correspond to the hidden rule state, a hallmark of model-based representation^57^. This compositional coding allows for a rapid transition of the population state-space when a rule change is detected. On the other hand, these same neurons display a prolonged update of stimulus-outcome associations following contingency changes, characteristic of model-free outcome-based learning^26^. This hybrid representation suggests that PPC_AC_ does not merely switch between static maps. Rather, it uses a model-based framework to guide rapid adaptation while simultaneously utilizing model-free feedback to fine-tune the new contingency. This dual-coding scheme enables the PPC to bridge the gap between abstract rule inference and the gradual acquisition of new sensory-motor associations.

Our findings extend the functional roles of the AC and IC beyond traditional sensory processing. While AC activity was dispensable for stable discrimination under familiar rules, it was essential for adaptive learning following contingency changes. This result aligns with evidence suggesting the auditory cortex is selectively recruited during the acquisition of new sensory-motor associations^32,38^. In contrast, the IC remained necessary for behavioral responses to cues across all task phases, consistent with its emerging role in integrating auditory and motor-related signals to support goal-directed behavior^47,48,58,59^. Extending this framework, our data suggest that PPC_IC_ projections convey action execution signals that stabilize sensory-to-motor transformation, whereas PPC_AC_ projections enable flexible, rule-dependent remapping. This division of labor may reflect an evolutionarily conserved strategy that prioritizes reliable execution while preserving the capacity for rapid adaptation. Moreover, imbalances within this dual-control system may lead to cognitive dysfunction.

Insufficient flexible updating or impaired stabilization could contribute to pathological perseveration or instability, respectively, as observed in various neuropsychiatric disorders^5–10,60–63^. Future studies in disease models will be essential to explore whether selectively rebalancing these parallel PPC outputs can restore adaptive behavioral control.

## Methods

### Animals

All experimental procedures were approved by the KAIST Institutional Animal Care and Use Committee (IACUC-18-100, IACUC-18-235). Animals were housed under controlled environmental conditions with a 12-hour light/dark cycle (8 am - 8 pm / 8 pm - 8 am) and provided *ad libitum* access to food and water. Throughout the experimental period, mice were individually housed at a controlled temperature (20 - 22[) and humidity (30 to 50%). We used C57BL/6 wild-type mice and B6.Cg-Igs7tm148.1(tetO-GCaMP6f, CAG-tTA2)Hze/J mice (Ai148, Jackson Laboratory, stock no. 030328) of both male and female, aged 2-6 months. Detailed information regarding the number of animals used in each experimental condition is provided in the corresponding figure legends and supplementary data tables.

### Surgery

#### Common procedure for animal surgery

Adult mice (P45-P80) were anesthetized with 1.5-2% isoflurane in oxygen, and their heads were fixed in a stereotaxic apparatus using ear bars. Body temperature was maintained at 37°C using a feedback-controlled heating pad (CWE Inc.). Following hair removal, the scalp was sterilized with 70% ethanol and povidone-iodine solution. A midline incision was made using surgical scissors for virus injection, or a portion of the scalp was removed for head plate implantation. After removing the periosteum above the skull, the exposed skull surface was cleaned with 70% ethanol, povidone-iodine solution, and sterile filtered phosphate-buffered saline.

#### Virus injection for dual-color retrograde tracing

For dual-color retrograde tracing, we injected ∼ 0.5 μl of retrograde adeno-associated virus (AAVrg) expressing tdTomato (pAAV-CAG-tdTomato, Cat# 59462-AAVrg, Addgene) and eGFP (pAAV-CAG-GFP, Cat# 37825-AAVrg, Addgene) into the right AC (posterior 2.5 mm and lateral 4.2 mm from bregma; depth 0.7 mm) and the right IC (posterior 5.2 mm and lateral 0.8 mm from bregma; depth 0.8 mm), respectively, of a wild-type mouse using a KDS Legato 130 Syringe Pump (RWD) at a rate of 0.3nl/sec. After 14 ∼ 15 days post-injection, brain samples were collected for histological analysis following transcardial perfusion.

#### Head plate implantation surgery for mouse head-fixation

For head plate implantation, mice were prepared as described above, including scalp incision and removal of connective tissues. We marked the PPC (posterior + 1.95 mm and lateral 1.4∼1.6 mm from bregma; depth 0.4 mm), AC, and IC for subsequent pharmacological inactivation experiments, *in vivo* extracellular recordings, and optogenetic manipulations. For lens implantation, a 1 mm × 1 mm area was demarcated on the skull 0.2 mm away from the PPC target coordinates. The custom-designed head plate was securely attached to the skull using a combination of small screws (Small Parts), cyanoacrylate glue (Loctite super glue 401, Henkel), and dental cement (Super-Bond C&B Kit, Sun Medical Co.). Following surgery, mice were closely monitored for their health status, and behavioral training began after a minimum recovery period of one week.

#### Surgical procedures for in vivo Ca^2+^ imaging with a miniature microscope

For circuit-specific Ca^2+^ imaging, ∼0.5 μl of AAVrg-hsyn-cre (pENN.AAV.hSyn.Cre. WPRE.hGH, Cat# 105553-AAVrg, Addgene) was injected into either the AC or IC of the right hemisphere in Ai148 mice before head-plate mounting. The exposed brain tissue at the injection site was covered with cured Kwik-Sil (WPI). Following a 7 – 10 day viral incubation period, mice began water restriction for behavioral training. During training on the auditory reversal-learning task (see Methods; Behavior), mice had full access to water until they returned to their baseline weight, ensuring proper health before the subsequent lens implantation surgery.

Gradient refractive index (GRIN) lens implantation was performed following established protocols^64^. Mice received subcutaneous injections of carprofen (5 mg/kg) and dexamethasone (5 mg/kg) at least 1 hour before surgery. Under anesthesia, a 1 mm × 1 mm craniotomy was made 0.2-0.3 mm anterior to the target field of view (FOV: PPC_AC_, AP -1.65 mm; PPC_IC_, AP -1.45 mm). The craniotomy location accounted for the working distance of the miniature microscope. A small linear incision was made in the cortical surface using a surgical knife (Cat# 10055-12, Fine Science Tools,) at the planned lens insertion site. The prism-attached GRIN lens (Cat# 1050-004624, Inscopix) was slowly inserted at a rate of 0.1 mm/20 s to a depth of 0.9-1.0 mm along the medio-lateral axis. The exposed brain tissue between the GRIN lens and skull was sealed with cured Kwik-Sil, followed by Super-Bond application to secure the lens. To protect the GRIN lens surface from debris during the recovery period, Kwik-Sil was applied to the top of the lens. Post-operative care included subcutaneous carprofen and dexamethasone injections for three consecutive days to manage pain and inflammation. After a 2-week recovery, water restriction was reinstated to resume behavioral training. Once mice regained stable performance (>0.75 Correct rate) in the auditory reversal task, the FOV for each mouse was examined for clear detection of neuronal activity. Following confirmation of a suitable imaging quality of FOV, a baseplate (Cat# 1050-004683, Inscopix) was attached to the GRIN lens using dental acrylic (Ortho-Jet, Lang Dental) and Super-Bond, then secured with a baseplate cover (Cat# 1050-004639, Inscopix).

#### Surgical procedure for optogenetic manipulation in task-performing mice

For circuit-specific optogenetic manipulations, AAVrg-CaMKIIα-ArchT (pAAV-CaMKIIα-ArchT-GFP, Cat# 99039-AAVrg, Addgene) was bilaterally injected into the AC or IC of wild-type mice in the test group. Control animals were injected with AAV2-CaMKIIα-eYFP (UNC) bilaterally. All viral injections were performed on the same day prior to head-plate implantation. After a 7 - 10 day period for virus expression, mice began water restriction for behavioral training. Once mice reached expert performance in the auditory reversal-learning task (see Methods; Behavior), they were provided unrestricted water access for at least 2 days to prepare for optic fiber implantation. Under anesthesia with 1.5 ∼ 2 % isoflurane, bilateral craniotomies (∼0.5 mm diameter) were made over the PPC using either manual drilling or a biopsy punch (Cat# 12-460-414, Fisher scientific, manufacturer). Optic fibers (200 μm core diameter, Cat# FT200UMT, Thorlabs) with ferrules (CFLC230-10, Thorlabs) were implanted into the PPC at a depth of 0.35mm and secured with Kwik-Sil (WPI) and Super-Bond. Following a 2-day recovery period, water restriction was resumed for optogenetic manipulation experiments.

### Behavior assay

#### Hardware setup

Auditory stimuli were digitally generated using custom MATLAB code (sampling rate 192,000 Hz, 16bit) with 5 ms sinusoidal ramps at stimulus onset and offset. A speaker (Ultrasonic Dynamic Speaker Vifa, Avisoft-Bioacoustics) was positioned 10 ∼ 12cm from the head-fixation apparatus. The sound pressure level (SPL) of both stimuli was calibrated to 75dB using a Reference Signal Generator and an externally polarized condenser microphone (CM16/CMPA, Avisoft-Bioacoustics), positioned at the same distance as the mouse’s left ear relative to the speaker. A custom-built lickometer was used to detect individual licking events during the task. Both outcomes (water reward and air puff punishment) were delivered through separate solenoid valves (EV-2-24, Clippard). All hardware components were controlled by Presentation (Neurobehavioral Systems) using a USB data acquisition board (USB-201, Measurement Computing).

#### Behavior training for the auditory reversal-learning task

Before behavioral training, mice underwent initial water deprivation until their body weight dropped below 80% of their pre-deprivation weight, typically after three days of deprivation. Body weight was monitored daily and maintained above 75% of the initial weight throughout the water restriction period. During behavioral experiments, mice were kept on ∼1 ml of water per day to ensure consistent performance. Task performance was evaluated based on hit rate, False Alarm (FA) rate, and Correct rate, which were calculated as follows:

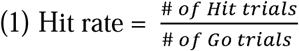

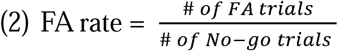

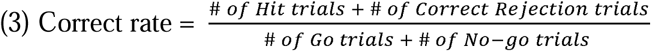

To assess perceptual flexibility in mice, we adapted and modified a reversal-learning task from previous research^23,65^. The task consisted of five stages:

1. Reward conditioning stage: Water-restricted mice received 1μl of water upon licking the water port. If mice did not initiate licking behavior within 3 minutes from the session start, water was manually delivered through the port until voluntary licking was established. Mice progressed to the next stage if they achieved more than 1,000 licks within 20 minutes.
2. R1 conditioning stage: Mice were trained to discriminate between Go (5kHz) and No-go (10kHz) pure tones (500 ms duration), presented pseudo-randomly in a 1:1 ratio. A 2-second response window followed a 500 ms delay after stimulus presentation. During Go trials, licking within the response window (hit) triggered a 3 μl water reward and a 2-second consumption period. If mice did not lick (miss trials), water was automatically delivered after the response window to encourage the Go stimulus-reward association. No-go trials had no consequences with licking (FA) or not licking (CR). The session included a 4-second inter-trial interval and terminated after 20 consecutive miss trials. Mice advanced to the next stage if they achieved a hit rate above 0.8.
3. R1 learning stage: Mice were trained to discriminate between 5 kHz and 10 kHz tones in a Go/No-go paradigm with reward and punishment. The paradigm was similar to the conditioning stage, except for two critical differences: no water reward for miss trials and negative outcomes for FA trials (4 - 8s timeout and/or 100 - 800 ms air puff).
4. Reversal-learning task with conditioning stage: Once mice reached a ≥0.75 Correct rate in the initial rule (R1), they proceeded to the reversal-learning stage, where the Go/No-go contingencies were reversed within a single session. To facilitate the rule transition, classical conditioning (CS) trials were introduced immediately following the reversal, during which a reward was delivered after Go stimuli regardless of licking behavior. CS trials were progressively reduced in 20-trial increments once mice achieved ≥0.75 Correct rates for both R1 and R2. When the number of CS trials reached 40, the delay period was extended from 500 ms to 1,000 ms. Mice exhibiting persistent cross-rule impulsive licking were returned to R1 training. Task performance during CS trials was evaluated based on anticipatory licking behavior^65^, with licks during the delay period classified as hits or FAs depending on the stimulus contingency. All subsequent experiments, including electrophysiology, drug treatments (MUS and PBS), Ca^2+^ imaging and optogenetics were conducted at this stage.
5. Reversal-learning expert stage: Mice that demonstrated a ≥0.75 Correct rate for at least three consecutive days in the R2 stage (without CS trials) were classified as reversal experts. Subsequent experiments, including drug treatments, Ca^2+^ imaging, and optogenetics were conducted at this stage.

#### Quantification of trial-number-to-reversal (TTR) and transition (TR) phase

To quantify the speed of adaptation to reversed contingencies (R2), we defined a metric termed trial-number-to-reversal (TTR). Following rule reversal, a 10-trial moving average of binary choice accuracy (Correct = 1, Incorrect = 0) was computed. TTR was defined as the final trial in the first sequence of three consecutive trials where this moving average exceeded 0.7. The TR phase was defined as the period between the contingency reversal and TTR (Figure 1). In sessions that included CS trials, TTR was calculated using both CS and standard trials, with anticipatory licking responses used to assess response accuracy.

#### Pharmacological inactivation

To investigate the role of specific brain regions in auditory reversal learning, we performed local neural inactivation through bilateral microinjections of either the GABA_A_ receptor agonist muscimol (MUS, 1 μg/μl; Sigma-Aldrich) or 1X PBS (control) into PPC, AC, and IC. Each hemisphere received 0.23 μl of solution. Behavioral testing in the auditory reversal task commenced 20 minutes post-injection to ensure adequate drug diffusion and effect. To control for individual variability and enable within-subject comparisons, mice were tested across multiple sessions in the following sequence: pre-injection baseline (no injection; Pre), muscimol injection (MUS), and PBS injection. Fluoro-ruby (Cat# AG335, Sigma-Aldrich) was injected with PBS to verify the injection site by histology. For specific subgroups (AC, N=1; IC, N=3; PPC, N=2), performance measures were averaged across sessions to generate a single data point per animal.

#### Optogenetic manipulations

For circuit-specific optogenetic inactivation during the reversal task, AAVrg-CaMKIIa-ArchT-GFP was bilaterally injected into either the AC (PPC_AC_) or IC (PPC_IC_), followed by bilateral optical fiber implantation in the PPC. To probe the temporal specificity of PPC inputs during behavioral adaptation, we employed two laser stimulation protocols (Type I and II inactivation):

1. Type I: Continuous green laser stimulation (532 nm, 10-12 mW) from 1 s before stimulus onset to the end of the response window (trial epoch).
2. Type II (Control): Trial-epoch stimulation delivered in pseudo-randomly selected trials.

For Type I, the number of stimulated trials was set by rounding up the TTR measured during matched laser-off sessions (up to a maximum of 80 trials) to ensure coverage of the TR phase. Type II served as controls to confirm that observed effects were specific to the transition stage.

#### *In vivo* electrophysiological recordings

### Neural activity recording during task

Neural recordings from AC and IC in task-performing mice were conducted using a 64-channel probe (P64-12, Diagnostic Biochips. Inc.). A 1 mm diameter craniotomy was made one day prior to recording and covered with an artificial dura-like material (Duragel, Cambridge Neurotech) to protect the exposed brain tissue. For each animal, 3 to 5 recording tracks were made in each target area. Neural data were acquired using an OpenEphys data acquisition board and software at 30kHz, with an amplifier gain of 192 for each channel. Signals were bandpass filtered with cutoff frequencies at 300 Hz and 6000 Hz. To confirm the recording sites and perform post hoc histological analysis, electrodes were coated with lipophilic dye (1,1’- dioctadecyl-3,3,303’- tetramethylindocarbocyanine perchlorate (DiI), Invitrogen) before insertion.

### Spike sorting for single-unit isolation

Spike sorting was performed using an automated spike-sorting MATLAB toolbox Kilosort to cluster units from raw data^66^. The resulting spike clusters were manually curated using Phy (Kwik team). Further quality control was conducted with custom MATLAB codes to ensure high-quality single-unit data. A total of 579 single units were recorded from AC and IC in 4 animals. To plot firing rate changes in representative neurons, peri-stimulus time histograms (PSTHs) of spike trains for 5kHz and 10kHz stimuli in R1 and R2 were generated separately by averaging trial-by-trial firing rates in 10-ms bins and smoothing with a Gaussian kernel (σ = 50 ms).

### Gain modulation index

For each stimulus-responsive neuron (Wilcoxon signed-rank test, p < 0.01), the change in firing rate during the stimulus period (0 to 500 ms) relative to the pre-stimulus baseline (-1500 to -500 ms) was computed on a trial-averaged basis, yielding Δ firing rates. For each stimulus frequency (5 kHz and 10 kHz, analyzed separately), the gain modulation index (GMI) was calculated by comparing response magnitudes between task phases (R2 or TR vs. R1), normalized by the maximum firing rate across these phases using a soft normalization parameter (a = 0.01).

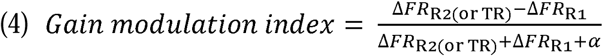

### Selectivity index

Neuronal selectivity for stimulus identity was quantified using receiver operating characteristic (ROC) analysis, as previously described^23,67^. Analyses were performed separately for R1 and R2 by comparing time-averaged firing-rate changes evoked by the 5kHz and 10kHz stimuli across trials. To capture both increased and decreased responses, firing-rate changes (Δfiring rates) were calculated for each trial as the difference between activity during the stimulus period (0 to 500 ms) and the pre-stimulus baseline (-1500 to - 500 ms). To minimize contamination from motor-related activity, trials with licking events occurring near the stimulus period (-500 to +750 ms relative to stimulus onset) were excluded from analysis. The stimulus selectivity index (SI) was then computed for each neuron from the area under the ROC curve (AUC) as follows:

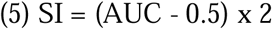

An SI value of +1 indicates consistently greater firing-rate changes in response to the 5kHz tone than the 10kHz tone, whereas an SI value of -1 indicates a preference for the 10kHz tone. Statistical significance of SI was assessed using a permutation test across trials (1,000 iterations; p < 0.01). To quantify changes in stimulus discriminability across task phases, we calculated ΔSI as follows:

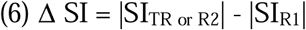

Neurons that did not exhibit a significant SI in any task phase were excluded from this analysis.

Outcome selectivity was quantified using the same ROC-based approach by comparing hit and CR trials during the response window (1500 to 3500 ms). In this case, an SI value of +1 indicates consistently greater firing-rate changes during hit trials, whereas −1 indicates greater responses during CR trials.

### Circuit-specific Ca^2+^ imaging and data analysis

#### Data acquisition and processing

We performed *in vivo* Ca^2+^ imaging using a miniature fluorescence microscope and the nVista 3.0 data acquisition system (Inscopix)^68,69^ with LED power of 0.2∼ 0.4 mW and a gain value of 1 ∼ 4, collecting data at 20 frames per second.

Data processing was performed using Inscopix Data Processing Software (IDPS). The preprocessing pipeline included spatial down-sampling (by 4), spatial filtering, and motion correction to mitigate motion artifacts. We used Constrained Non-negative Matrix Factorization for micro-endoscopic data (CNMF-E) to identify regions of interest (ROIs) corresponding to individual cells. ROI detection was configured with a cell diameter between 5 to 7 pixels and standard thresholds for minimum pixel correlation (0.60 to 0.80) and minimum peak-to-noise ratio (8.00 to 10.00) to ensure Ca^2^ signal detection while minimizing false positives. After automated ROI detection, we manually curated the results to match fluorescence traces (ΔF/F_0_) with ROIs and eliminate contamination from out-of-focus neuropil fluorescence. To correct for fluorophore photo-bleaching, we applied a correction method from a previously published protocol^70^, using the following equation:

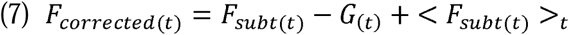

This equation calculated the corrected fluorescence signal (F_corrected(t)_) by subtracting slow baseline fluctuations averaged over a 300-second sliding window (G_(t)_) and adding back the overall mean fluorescence across the recording session (<F_sub(t)_>_t_).

#### Rastermap visualization

Out of 798 recorded neurons (594 PPC_AC_ neurons and 194 PPC_IC_ neurons), we selected neurons with an average ΔF/F_0_ greater than 0.05 (283 PPC_AC_ neurons and 111 PPC_IC_ neurons). To visualize task-related Ca^2+^ activity, we applied Rastermap^52,53^, a k-means clustering-based sorting algorithm that highlights temporally correlated patterns in neural population activity. Among the tunable parameters, we selected a cluster number of 50 for sorting population activity. Importantly, choosing alternative values (e.g., 25 clusters) did not alter the overall patterns or conclusions of our observations (Data not shown).

#### Principal Component Analysis (PCA)

Ca^2+^ activity was averaged for each trial type (hit and CR) and task phase (R1, TR, and R2), then concatenated to form a 6T x N matrix, where T represents the duration of each trial segment (2 second pre-stimulus onset and 6 seconds post-stimulus onset) and N is the number of imaged neurons. We then performed principal component analysis (PCA) using pca function in Matlab library). Neural trajectories were computed by projecting the data onto the first three principal components (PCs).

#### Subspace overlap analysis

Normalized Ca^2+^ traces were trial-averaged in 50-trial segments S_i_ (i = 1, … 7) across R1 (S1-S3; 1 to 150 trials), TR (S4; R2 to TTR + 15 trials) and R2 (S5-S8; 1 to 150 trials post S4). We then performed PCA using *scikit-learn* (Python library) to compute low-dimensional neural subspaces for each segment. To quantify subspace overlap, we calculated the cosine of the principal angle between each pair of subspaces, as previously published^71^. The principal angle (*θ_p_*) between two subspaces *U* and *V* corresponds to the largest angle between any two pairs of vectors in *U* and *V*, providing a measure of their alignment. For each segment *S_i_*, we extracted the first N principal components and organized them into an N × N matrix **R(***S_i_*), defining the subspace of that segment. The cosine of the principal angle between subspaces *S_i_* and *S_j_* was computed using the method described by Knyazev and Argentati^72^, as follows:

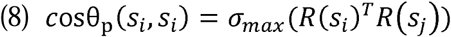

Where *σ_max_*(A) denotes the largest singular value of matrix A.

#### Generalized linear models (GLM)

We used generalized linear models (GLM) to regress recorded Ca^2+^ signals against task event variables^70,73,74^. To capture contingency-dependent neural dynamics, GLMs were trained on specific task phase (R1, TR or R2) rather than on the full session. Ca^2+^ responses for each neuron were modeled as a linear sum of task components and events, including lick-bout onset and offset, Go stimulus (10kHz) and No-go stimulus (5kHz), hit, FA, CR, and miss.

The neural response at time t (*y_t_*) was modeled by the sum of the bias term (*β_0_*) and the weighted sum of various binary predictors (*x_i_*) as follows:

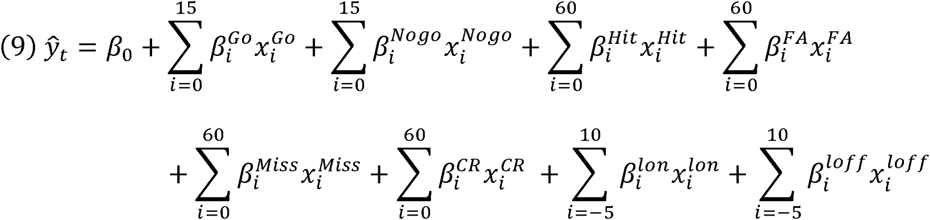

Predictors for each task variable were convolved with temporal filters to capture the slow dynamics of Ca^2+^ signals^75^. Predictors for Go 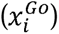 and No-go 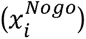 stimuli indicated stimulus presence, while lick predictors represented onset 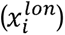 and offset 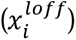 of lick bouts, defined as sequences with inter-lick intervals of less than 2 seconds. Predictors for behavioral outcomes, including hit 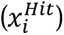, FA 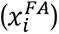, CR 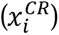, miss 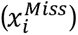, included time lags to cover the duration of stimulus presentation, reward consumption, and slow Ca^2+^ dynamics. Time lags were set to 0-1.5 second for stimulus predictors, 0-6 seconds for outcome predictors, and -0.5 to 1 s for lick-related predictors, where 0 denotes the stimulus (for stimulus and outcome) or lick onset time. The full model contained up to 321 coefficients, including the bias term (*β_0_*).

To avoid overfitting, we used a bottom-up model selection approach. We first fit single-variable GLMs for each neuron using *ElasticNet* linear regression (scikit-learn, Python library), which combines L1(lasso) and L2 (ridge) regularization. Model performance was evaluated using twenty-fold cross-validation, with 30% of trials held out for validation. For each neuron, the task variable (TV) whose model produced the highest mean cross-validated R^2^ score significantly greater than 0 (Student’s t-test) was selected as the initial predictor. Neurons for which no TV reached significance were classified as not fitted by the GLM. Additional TVs were then incorporated iteratively. At each step, we compared model fits with and without the added TV and retained the new TV only if it significantly improved model performance (two-sample Kolmogorov–Smirnov test). This process was repeated until no further improvement was detected. The ElasticNet regularization parameter α was optimized across a broad range (1 x 10^-5^ to 10) to maximize model performance.

To analyze patterns of neuronal multiplexing (co-encoding of multiple TVs), we identified all unique combinations of co-encoded variables and ranked multiplexing patterns by their frequency across neurons. For neurons encoding more than two TVs, all pairwise combinations were counted separately. For example, a neuron encoding “Hit,” “FA,” and “CR” contributed one count each to the pairs “Hit-FA”, “Hit-CR”, and “FA-CR”. This approach enabled us to quantify how frequently each pair of TVs was co-encoded across the population. Co-encoding relationships were visualized using a customized circular Sankey diagram (Matlab File Exchange, ID: 128679)^76^. In these diagrams, curved links connect pairs of co-encoded variables, and the thickness of each link is proportional to the frequency of co-occurrence.

### Histology and imaging

#### Fluorescence imaging of brain sections

Histological experiments were conducted to verify anatomical circuit tracings, confirm targeting sites after pharmacological inactivation, identify electrode tracks following *in vivo* recordings, and validate optogenetic manipulation and Ca^2+^ imaging sites. Mice were deeply anesthetized with isoflurane and intraperitoneal injection of avertin (2,2,2,-Tribromoethanol; 125-250 mg/kg; Sigma-Aldrich), and transcardially perfused with 10ml of phosphate-buffered saline (PBS) followed by 4% paraformaldehyde (wt/vol in 1X PBS). Brain samples were carefully extracted and post-fixed for 4 hours at room temperature, then washed 3 times in 1X PBS for 10 minutes each. The samples were transferred to a filtered 30% sucrose solution (wt/vol in 1X PBS) and allowed to sink completely for 1∼2 days at 4°C. Brain samples were embedded in the optimal cutting temperature (OCT) compound (Tissue-Tek, Sakura Finetek), rapidly frozen at -80°C, and coronally sectioned at 30 μm (for tracing) or 40 μm (for site verification) using a cryostat (Leica). Sections were washed three times in 1X PBS for 10 minutes to remove residual OCT medium. The sections were mounted on glass slides with a mounting medium containing 4′,6-diamidino-2-phenylindole (DAPI, Vector Labs) for nuclear staining and tissue visualization. Fluorescent images were acquired using a slide scanner (Axio Scan Z1, Carl Zeiss).

#### Tracing analysis

For dual-retrograde tracing analysis, three coronal brain sections (30 µm thick) were collected from each of three mice. Sections were selected based on stereotaxic coordinates from the Paxinos atlas (AP -1.70 mm, -1.84 mm, and -2.06 mm from the bregma). Using IMARIS software (Oxford Instruments), brain section images were aligned to the reference atlas, and anatomical boundaries were matched to delineate the PPC, including its medial (PPC-M) and lateral (PPC-L) subdivisions. Labeled cells within these boundaries were detected using the IMARIS Cell Detection function, with DAPI colocalization used to confirm accurate cell identification. The spatial coordinates (x, y, z) of verified cells were exported for custom MATLAB-based analysis. To assess layer-specific distributions, atlas boundaries were imported into IMARIS to guide selection of DAPI signals corresponding to pia and ventricular surfaces. Cortical depth was normalized by regression fitting of DAPI signals along these boundaries, with the pia-to-ventricle distance scaled to a depth of 1. This normalization allowed direct comparison of relative laminar distributions between PPC_AC_ and PPC_IC_ populations.

#### Whole-brain clearing and 3D imaging

Whole-brain clearing was performed using the Binaree Tissue-Clearing Kit (HRTC-012 and HRMO-006, Binaree, Inc.) following a previously published protocol^77^. After perfusion and brain extraction, the tissue-clearing process proceeded as follows: The brain was incubated in a starting solution at 4°C for 24 hours. The brain was then transferred to tissue-clearing solutions and incubated at 37°C with 50 rpm agitation for 48 hours. This clearing step was repeated to achieve complete tissue transparency. The brain was rinsed three times in a washing solution, then incubated in a mounting and storage solution at 37°C for 24 hours. For imaging, a Lightsheet 7 fluorescence microscope (Carl Zeiss) was used to acquire z-stack images of the cleared brain. The resulting images were reconstructed in 3D using IMARIS software.

### Model Simulation

#### Neural network configuration

To simulate an auditory Go/No-go task with within-session rule reversal, we implemented a hierarchical feed-forward network consisting of sensory (input, IC, AC), modulatory (PPC_AC_, PPC_IC_) and action output modules (Figure 6). Auditory tone information was propagated from the IC to the AC and integrated to generate action logits corresponding to lick versus no-lick responses. Two parallel modulatory modules controlled the gain of the IC and AC layers. Specifically, PPC_AC_ monitored outcome contingency statistics (hit, miss, FA and CR), computed errors as deviations from the expected contingencies, and generated a signal that promoted adaptive updating in the AC. PPC_IC_ computed a goal-directed action signal (the difference between the probabilities of lick and no-lick responses) and modulated IC gain to stabilize action execution.

#### Reversal learning

Agents were trained on a Go/No-Go task with a within-session reversal of stimulus-action contingencies, identical to the behavioral experiment. Actions were sampled from a softmax policy, and scalar rewards were assigned according to the task rule: r = +1 for licking on Go trials and r = −1 for licking on No-go trials. Parameters were optimized using a standard policy-gradient objective, 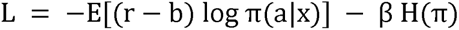, which included an entropy bonus (H(π): categorical entropy with β =1×10^−3^) to encourage exploration. The reward baseline b was computed as an exponential moving average of rewards, updated as b ← λ b + (1−λ)E[r] with λ=0.95.

#### Flexible-stable synaptic weight updates

Synaptic weights were updated using a flexibility-weighted meta-update rule^56,78^, in which each parameter’s gradient update was gated by its assigned flexibility. Stable pathways (input–IC, IC–readout) were assigned low flexibility, whereas IC–AC and AC–readout pathways were assigned higher flexibility, enabling selective adaptation following rule reversal. Low-flexibility synapses were anchored to their initial values, thereby suppressing updates as parameters deviated from this anchor.

Specifically, for each synapse, the deviation from the anchor weight was defined as Δ = *p* – *w_0_*, and parameters were updated according to

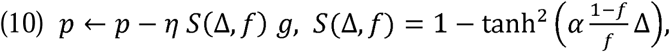

where *g* denotes the gradient of the loss, f ∈ [0,1] represents synaptic flexibility, with *a* = 4 controlling the sharpness of the gating function, chosen to provide a clear separation between flexible and stable update regimes. Under this rule, high-flexibility synapses (*f* ≈ 1) undergo near-standard gradient-based updates, whereas low-flexibility synapses (*f* ≪ 1) rapidly suppress updates as they deviate from the anchor, effectively preserving previously learned synaptic structure.

#### Agent performance and inactivation conditions

Circuit-specific inactivation (AC, IC, PPC_AC_ and PPC_IC_) was simulated by multiplicative suppression of activity or flexibility signals in the corresponding modules, matching the experimental manipulations. Behavioral performance was quantified by stimulus-conditioned lick probabilities, estimated from held-out trials, for hit and FA responses.

#### Alternative network architectures for comparison

To evaluate the functional advantage of the parallel circuit motif, we implemented an integrated PPC model and two control networks with uniform synaptic properties. In the integrated PPC model, a single population integrated both goal-directed and contingency-related errors into a global population error signal. Crucially, this global error signal was jointly coupled to both stability and flexibility modulation, preventing the independent control characteristic of the parallel model. All simulation scripts are available at https://github.com/seungheelee1789/PPC_RL_Jung.

### Statistical analysis

Data analysis was conducted using Microsoft Excel, custom MATLAB (MathWorks) and Python scripts. All data are presented as means ± standard error of the mean (SEM) unless otherwise specified. ’N’ denotes the number of mice, while ’n’ represents the number of sessions or neurons. Statistical differences were evaluated through repeated measures two-way ANOVA, Wilcoxon signed-rank test (non-parametric, paired datasets), Paired t-test (parametric, paired datasets), Wilcoxon rank-sum test (non-parametric, unpaired datasets), two-sided chi-square test (non-parametric, frequency counts), Mood’s median test (non-parametric, unpaired datasets), Sign-test (non-parametric, zero median of single population), and Two-sample Kolmogorov-Smirnov test. All the statistical significances are indicated as *, **, ***, **** for *p* < 0.05, *p* < 0.01, *p* < 0.001, and *p* < 0.0001, respectively.

## Acknowledgments

This work has been supported by grants to S.H.L. from the Institute for Basic Science (IBS-R002-A2) and to S.P. from the National Research Foundation of Korea (RS-2002-NR070602).

## Author Contributions

E.J. and S.L. conceptualized and initiated the project. E.J., J.L., and S.L. designed experiments and wrote the manuscript. E.J. and J.L. jointly performed data analysis and visualization. E.J. performed behavioral training, histology, Ca^2+^ imaging, optogenetic experiments, and some *in vivo* recordings. J.L. established *in vivo* electrophysiology recordings and implemented subspace overlap and GLM analyses. W.C. designed the computational model. S.P. provided advice and conceptual guidance. G.R. conducted some *in vivo* recordings and behavioral training. G.K. performed brain clearing and cell detection.

## Data and code availability

Source data are provided with this paper. All the representative data and code that support the findings are publicly available on GitHub: https://github.com/ seungheelee1789/PPC_RL_Jung. Further requests for the data generated in this study can be directed to the corresponding author.

## Ethics declarations

### Competing interests

The authors declare no competing financial interests.

## Extended Data Figures

**Supplementary Figure 1.**
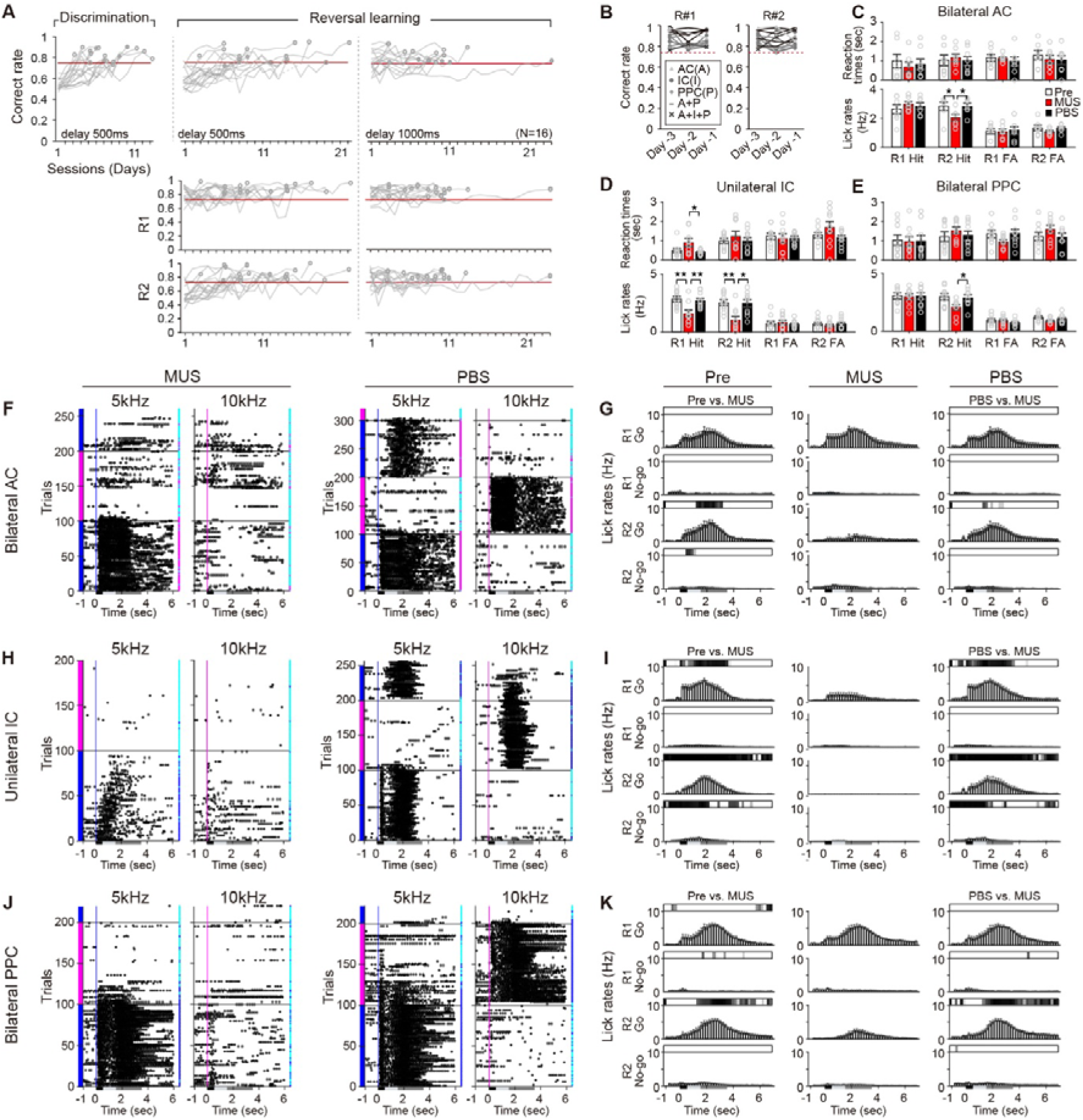
Task performance during within-session auditory reversal learning and effects of MUS inactivation. **A.** Performance changes in mice during auditory reversal learning. Top, Correct rates across training stages in mice used for single-unit recordings (N=3, blue) and MUS experiments (N=13, gray). Left, auditory discrimination (R1 only) with a 500 ms delay of the response window from the stimulus offset; middle, reversal learning with a 500 ms delay; right, reversal learning with a 1000 ms delay. Red horizontal lines indicate the expert-level performance threshold (Correct rate ≥ 0.75). Bottom, Correct rates for R1 and R2 trials during reversal learning with 500 ms (left) and 1000 ms (right) delays. **B.** Baseline performance prior to MUS inactivation. Mice reached Correct rates > 0.75 for both rules across three consecutive days before MUS injections into AC, IC, or PPC. **C.** Lick reaction times (Top) and lick rates (Bottom) for hit and FA trials during AC inactivation. Circles, individual sessions; open bars, Pre; red bars, MUS; black bars, PBS. Wilcoxon rank-sum test; R2 hit lick rates, Pre vs. MUS (p = 0.034); Mus vs. PBS (p = 0.024). **D.** Same as **C**, but for IC inactivation sessions. Wilcoxon rank-sum test; reaction times of R1 hit trials, MUS vs. PBS (p = 0.053); lick rates of R1 hit trials, Pre vs. MUS (p = 0.008); MUS vs. PBS (p = 0.010); lick rates of R2 hit trials, Pre vs. MUS (p = 0.004); MUS vs. PBS (p = 0.013). **E.** Same as **C**, but for PPC inactivation sessions. Wilcoxon rank-sum test; R2 hit lick rates, MUS vs. PBS (p = 0.019). **F.** Example lick rasters from AC inactivation sessions. Left, MUS; right, PBS. **G.** Session-averaged lick histograms for hit and FA trials during Pre (left), MUS (middle), and PBS (right) sessions targeting bilateral AC. Top bars, gray-scaled p-values relative to MUS sessions (Wilcoxon signed-rank test). **H-I.** Same as **F-G**, but for IC inactivation sessions **J-K.** Same as **F-G**, but for PPC inactivation sessions.

**Supplementary Figure 2.**
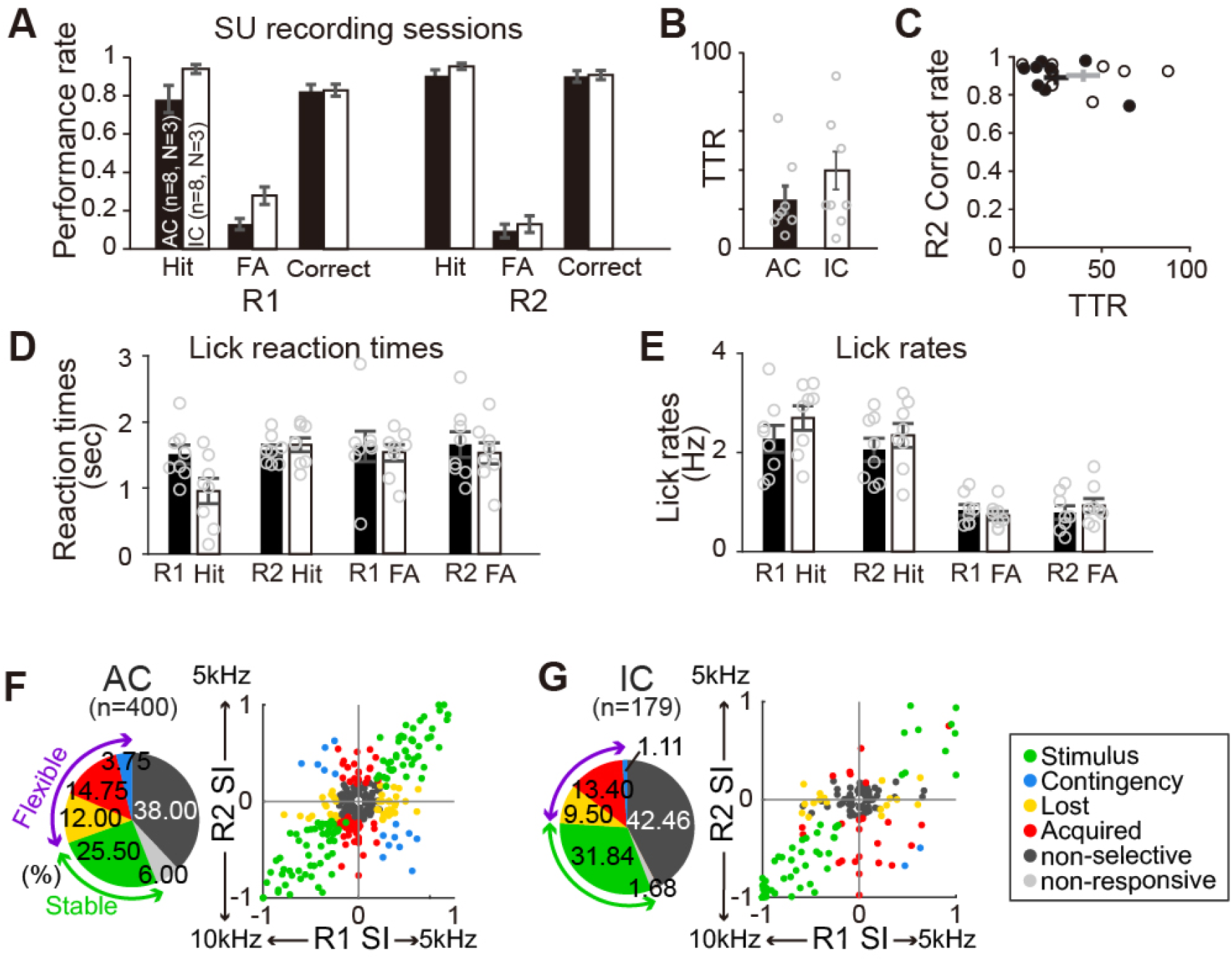
Behavioral performance during single unit recordings in AC and IC. **A.** Average hit, FA, and Correct rates in mice recorded from AC (N=3, n=8 sessions; black bars) and IC (N=3, n=9 sessions; open bars). No significant group differences (two-way ANOVA without repetition). **B.** Average TTRs for AC-recorded (24.37 ± 6.95) and IC-recorded (42.66 ± 10.16) sessions. Circles represent individual sessions. **C.** Scatter plot of TTRs vs. R2 Correct rate (AC = 0.87 ± 0.04; IC = 0.93 ± 0.01). Circles, individual sessions (black, AC; empty, IC); crosses, group means ± SEM. **D-E.** Lick reaction times (**D**) and lick rates (**E**) for hit and FA trials. Circles, sessions; black bars, AC-recording sessions; open bars, IC-recording sessions. No significant group differences (Wilcoxon rank-sum test). **F-G.** Left, pie charts showing fractions of neurons in each SI category in AC (**F**) and IC (**G**). Right, scatter plots of SI in R1 versus SI in R2 for individual neurons in AC (**F**) and IC (**G**). Inset, color key for six neuron categories defined by rule-dependent changes in SI: stimulus-selective (green), contingency-selective (blue), lost selectivity (yellow), acquired selectivity (red), non-selective (dark gray), and non-responsive (light gray).

**Supplementary Figure 3.**
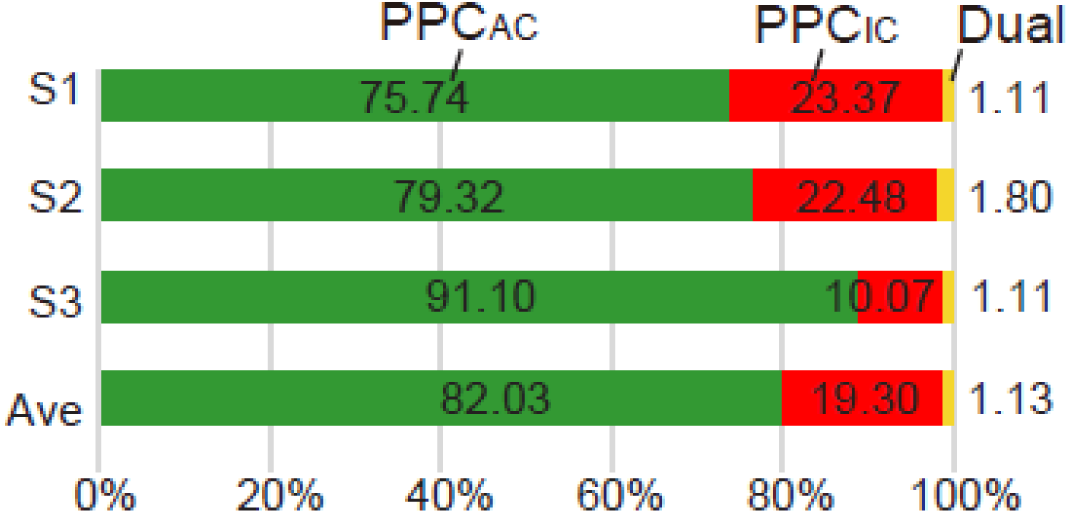
Quantification of projection-specific PPC neurons. Proportions (%) of PPC_AC_, PPC_IC_, and dual-projecting neurons across individual brain samples (S1-S3, N=3) and group average (bottomc).

**Supplementary Figure 4.**
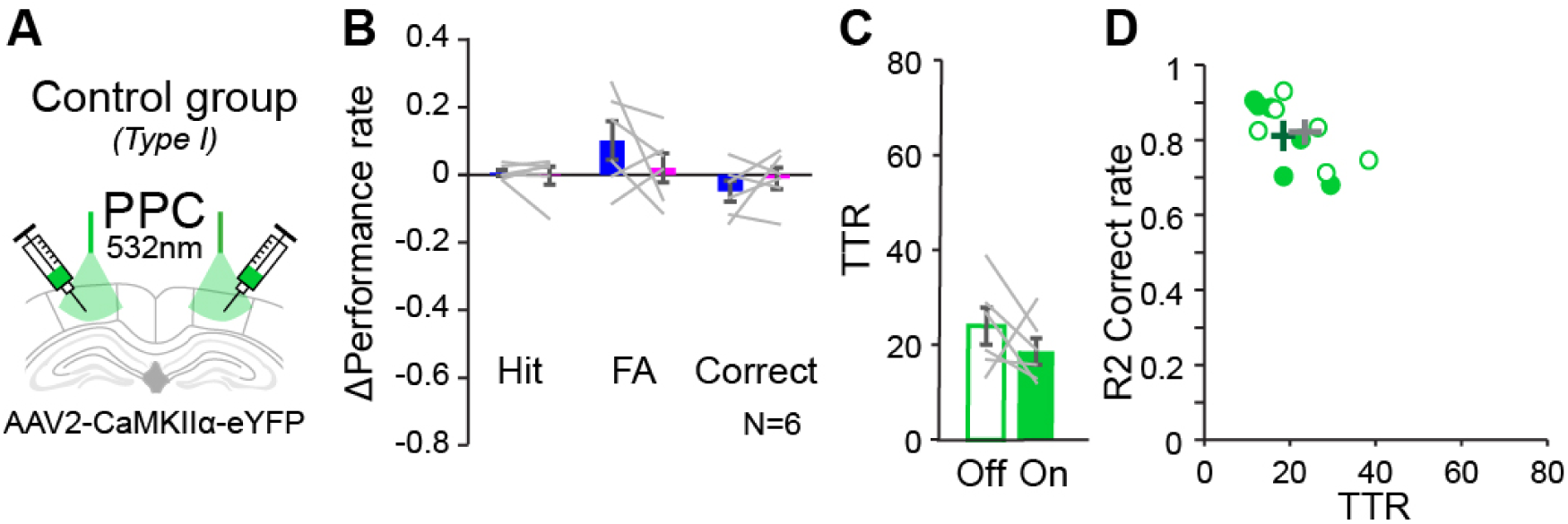
Control experiment for optogenetic manipulations of PPC_AC_ and PPC_IC_ circuits during auditory reversal learning. **A-D.** Control green-light (532 nm) stimulation of PPC during Type I optogenetic inhibition experiments. **B.** Δ Performance (light-on - light-off) for hit (R1, 0.006 ± 0.009; R2, -0.002 ± 0.026; n.s.), FA (R1 = 0.102 ± 0.057; R2 = 0.021 ± 0.043; n.s.), and Correct rates (R1, -0.049 ± 0.032; R2, -0.010 ± 0.032; n.s.). Wilcoxon signed-rank test. **C.** TTRs during light-off (24.00 ± 3.89) and light-on (18.83 ± 2.77) sessions; p = 0.009. Wilcoxon signed-rank test. **D.** Scatter plot of TTRs vs. R2 Correct rates. Green-open circles, light-off sessions; green-filled circles, light-on sessions; Crosses indicate group means ± SEM (gray, light-off; dark green, light-on)

**Supplementary Figure 5.**
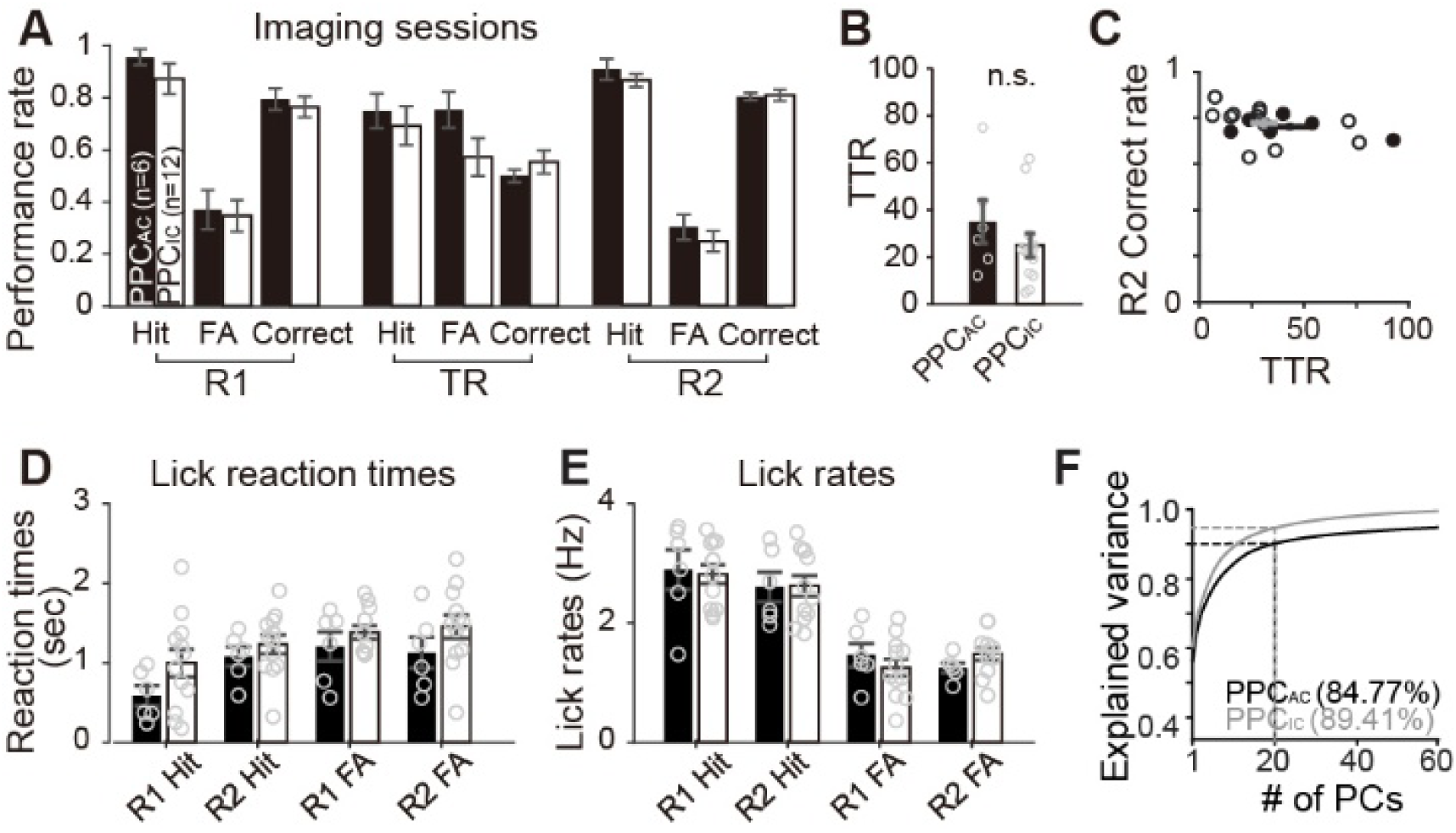
Behavioral performance during *in vivo* imaging of PPC_AC_ and PPC_IC_ neurons and robustness of neural population analyses. **A.** Average hit, FA, and Correct rates for PPC_AC_ (n=6, black bars) and PPC_IC_ (n=12, open bars) imaging groups. No group differences (two-way ANOVA without repetition). **B.** Average TTRs for PPC_AC_ (34.50 ± 9.02) and PPC_IC_ (24.58 ± 5.12). Circles, individual sessions. **C.** Scatter plot of TTRs vs. R2 Correct rates across all imaging sessions (PPC_AC,_ 0.76 ± 0.01; PPC_IC_, 0.77 ± 0.02). Circles, imaging sessions (black, PPC_AC_; empty, PPC_IC_); crosses, group means ± SEM (black, PPC_AC_; gray, PPC_IC_). **D-E.** Lick reaction times **(D)** and lick rates **(E)** for hit and FA trials. Circles, individual sessions; black bars, PPC_AC_; open bars, PPC_IC_. No significant group differences (Wilcoxon rank-sum test). **F**. Explained variance of Ca^2+^ population activity for PPC_AC_ (black) and PPC_IC_ (gray) neurons across task phases (same dataset as Figures 4E and **4F**). The top 20 PCs capture ∼80% of total variance, validating the use of 20 PCs.

**Supplementary Figure 6.**
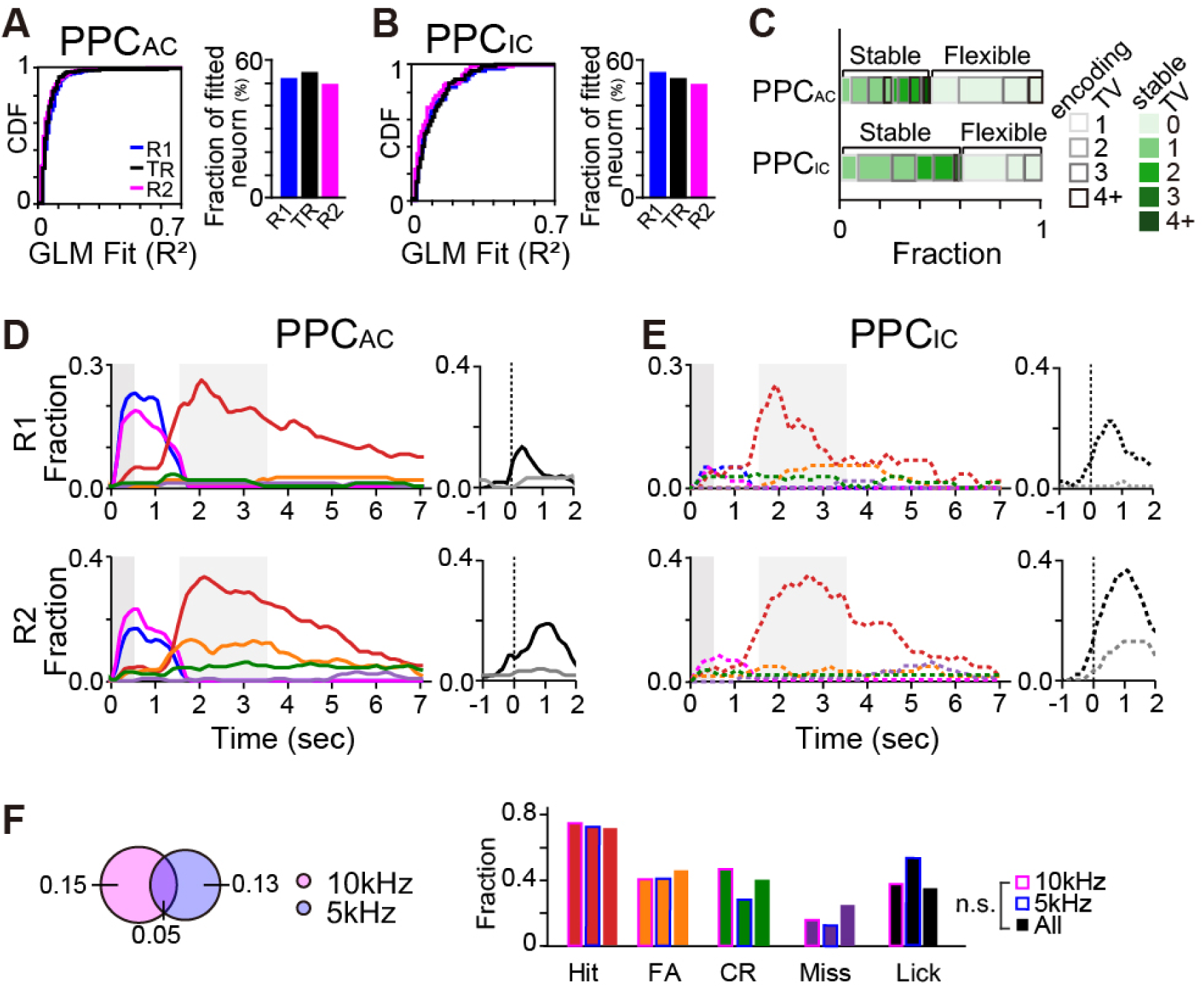
Temporal dynamics of task variable encoding in projection-specific PPC populations across rules. **A.** Left, cumulative distribution of GLM-fitted R^2^ values across rules. Blue line, R1; black line, TR; magenta line, R2. Right, fraction of neurons encoding each task variable across rules. Blue bar, R1; black bar, TR; magenta bar, R2. Noted that similar proportions of neurons were fit across phases. **B.** Same as **A**, but for PPC_IC_ neurons. **C.** Neurons grouped by the number of TVs encoded during TR (box outlines); filled colors indicate stability of TV encoding across rules. PPC_AC_ exhibited lower pattern stability (0.57) than PPC_IC_ (0.40), indicating more stable TV encoding in PPC_IC_ across rule changes. **D.** Left, time-resolved fractions of PPC_AC_ neurons encoding each task variable (color-coded) during R1 (top) and R2 (bottom), aligned to stimulus onset. Right, fractions encoding lick-bout onset (black) and offset (gray), aligned to their respective motor events. **E.** Same as **D**, but for PPC_IC_ neurons. **F.** Left, Euler diagrams of PPC_AC_ neurons encoding 10kHz and 5kHz stimuli. Right, comparison of co-encoding fractions among 10kHz-encoding (magenta outline), 5kHz-encoding (blue outline), and all TV-encoding (no outline) PPC_AC_ neurons. No significant difference (two-sided chi-square test).

**Supplementary Figure 7.**
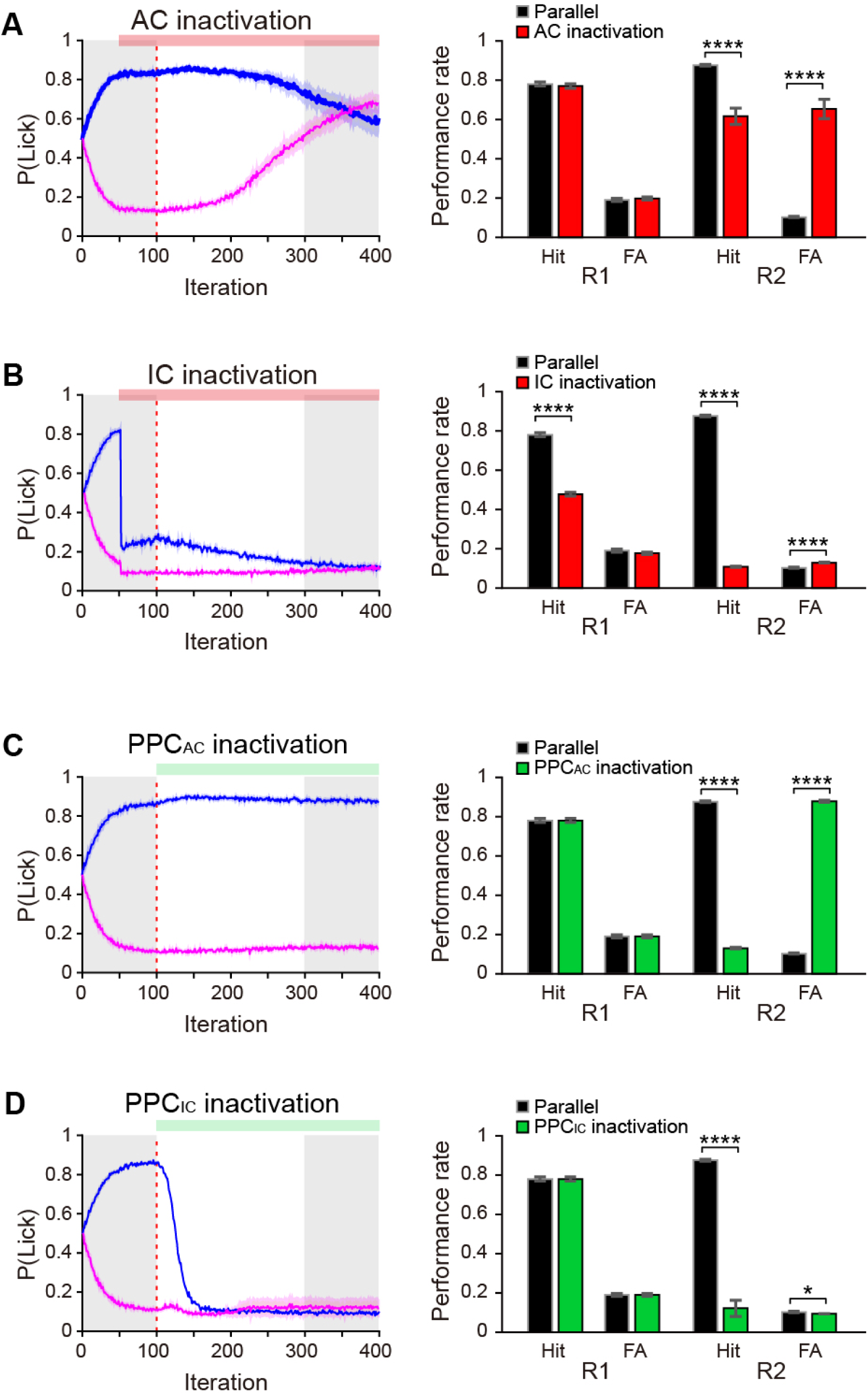
Inactivation simulations support a parallel top-down control model. **A.** Left, lick probability across learning iterations under simulated AC inactivation across task rules. Right, average hit and FA rates under control conditions (intact AC; open bars) and AC inactivation (red bars) across simulations (N = 20). Wilcoxon rank-sum test; R1 hit rate (n.s.); R1 FA rate (n.s.); R2 hit rate (p < 0.00001); R2 FA rate (p < 0.00001). **B.** Same as **A,** but for IC inactivation. Wilcoxon rank-sum tests; R1 hit rate (p < 0.00001); R1 FA rate (n.s.); R2 hit rate (p < 0.00001); R2 FA rate (p < 0.00001). **C.** Left, lick probability across learning iterations under simulated PPC_AC_ pathway inactivation following contingency reversal. Right, average hit and FA rates under control (circuit on; open bars) and PPC_AC_ inactivation (circuit off; green bars); N = 20 simulations. Wilcoxon rank-sum test; R1 hit rate (n.s.); R1 FA rate (n.s.); R2 hit rate (p < 0.00001); R2 FA rate (p < 0.00001). **D.** Same as **C**, but for PPC_IC_ pathway inactivation. Wilcoxon rank-sum test; R1 hit rate (n.s.); R1 FA rate (n.s.); R2 hit rate (p < 0.00001); R2 FA rate (p = 0.026).

## Supplementary Tables

**Supplementary Data Table 1.**
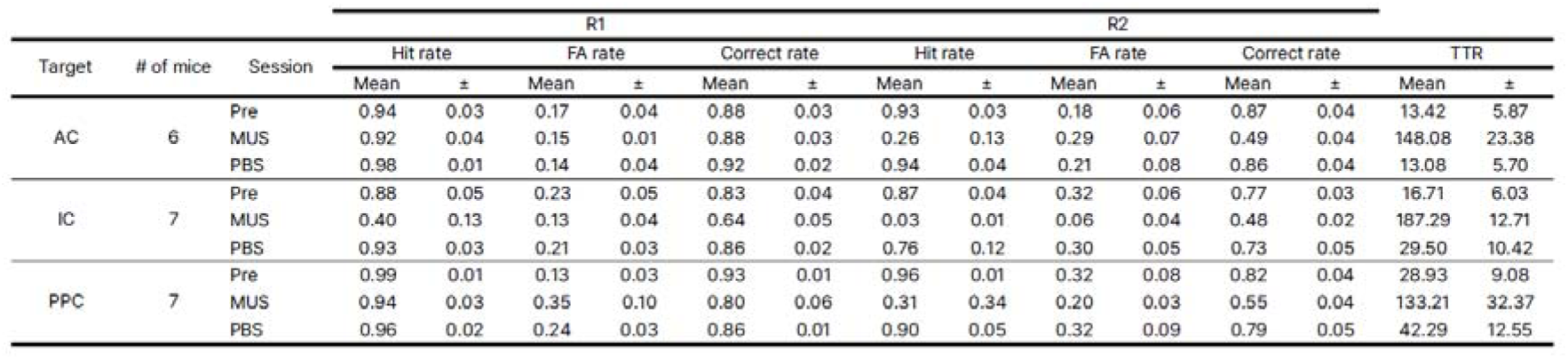
Behavior performance rates and TTRs of mice used for MUS experiments.

**Supplementary Data Table 2.**
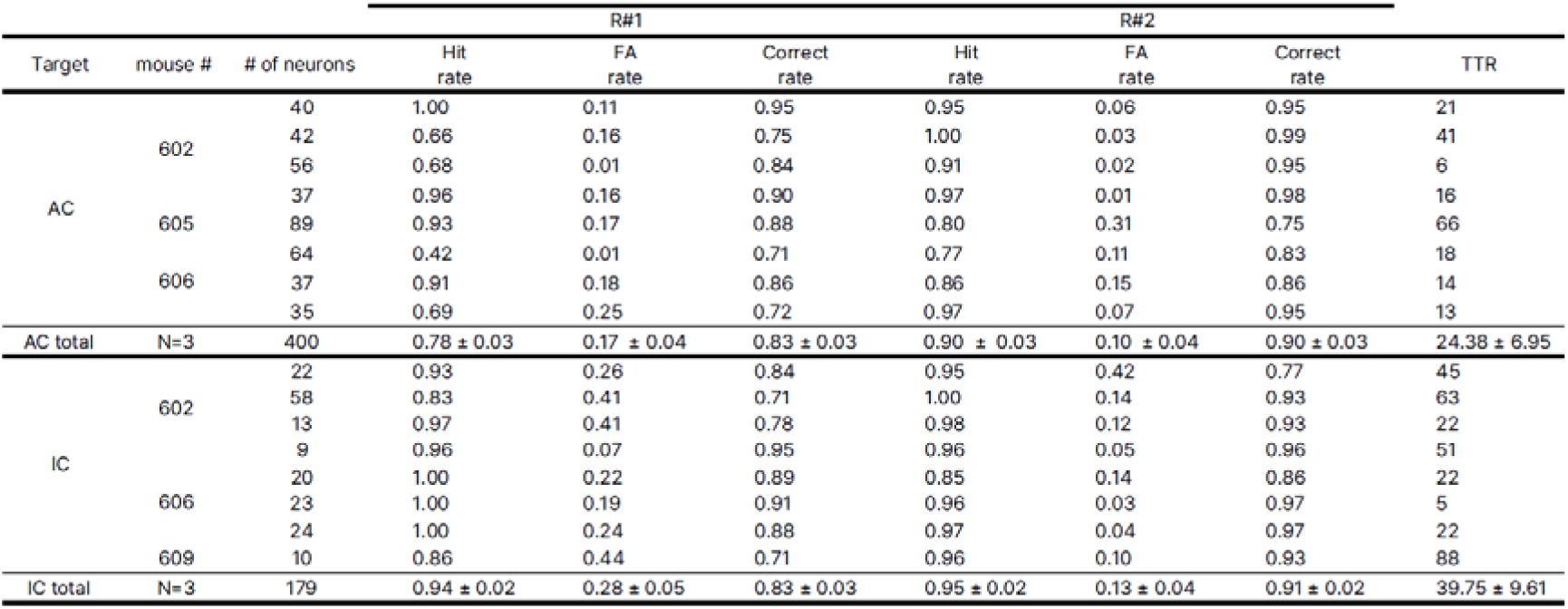
Behavior performance rates and TTRs of mice used for electrophysiology.

**Supplementary Data Table 3.**
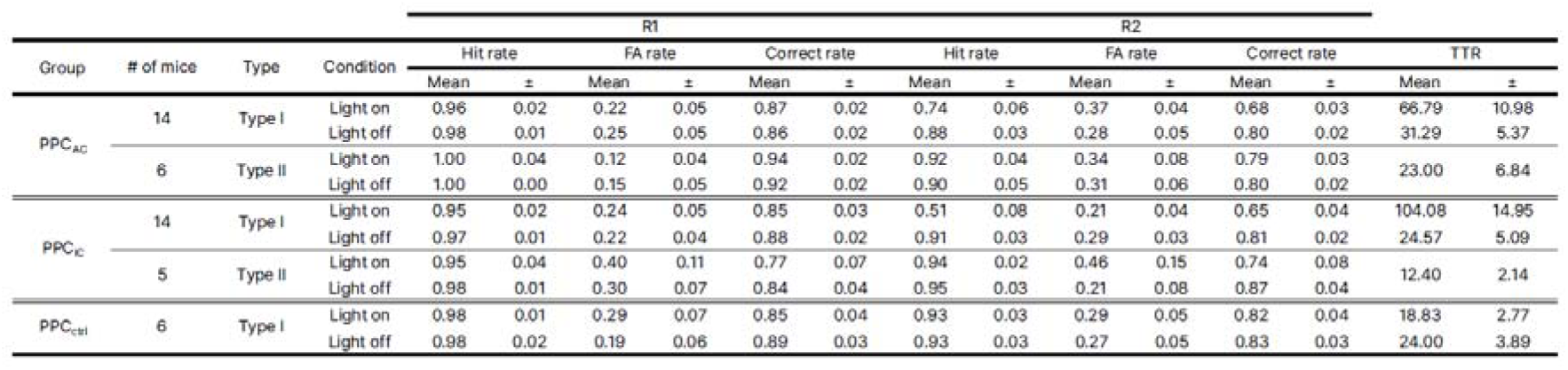
Behavior performance rates and TTRs of mice used for optogenetic manipulations.

**Supplementary Data Table 4.**
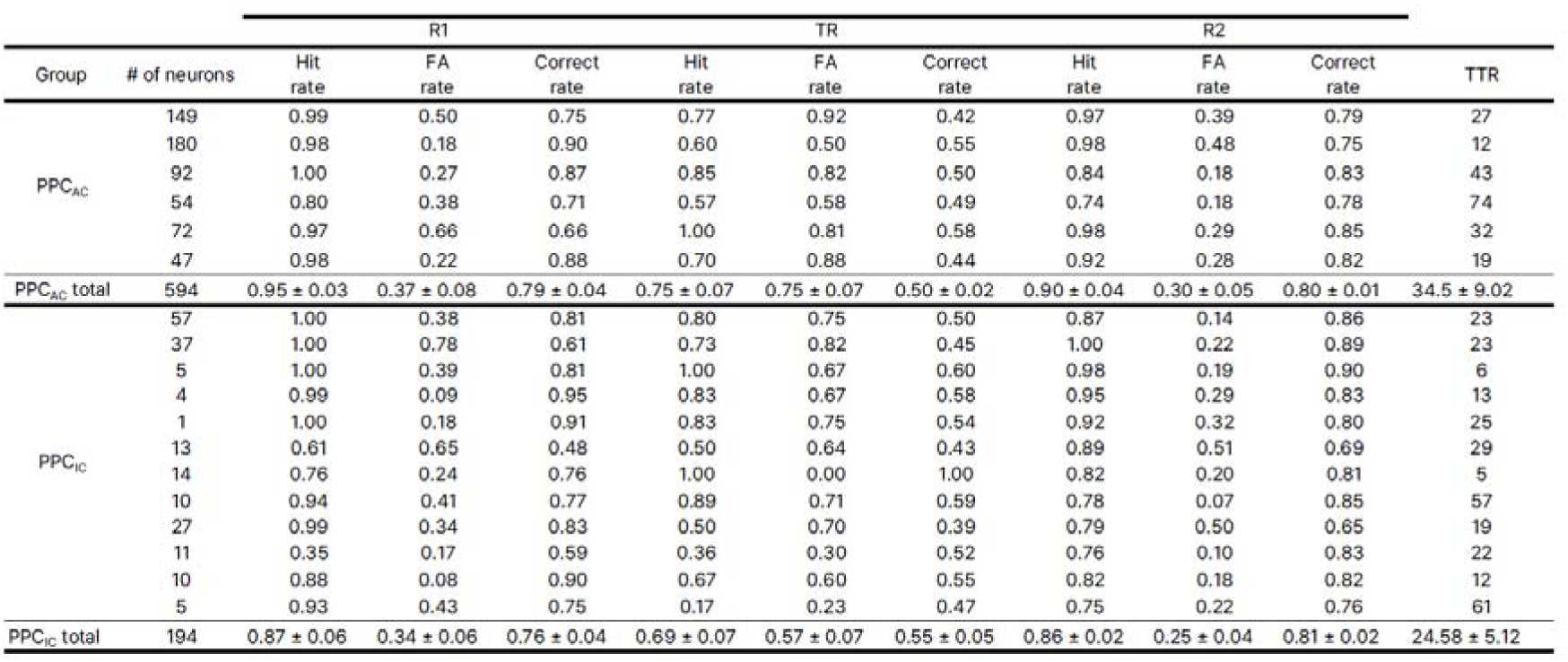
Behavior performance rates and TTRs of mice used for Ca^2+^ imaging.

## Notes

https://github.com/seungheelee1789/PPC_RL_Jung

### Competing Interest Statement

The authors have declared no competing interest.

### Summary of Updates

The revised manuscript now includes a feedforward network model as well as additional analyses.

