## Supplementary Tables for "Parallel circuits in the posterior parietal cortex balance behavioral flexibility and stability"

| Target | # of mice | Session | R1 | | | | | | R2 | | | | | | | |
|---|---|---|---|---|---|---|---|---|---|---|---|---|---|---|---|---|
| | | | Hit rate | | FA rate | | Correct rate | | Hit rate | | FA rate | | Correct rate | | TTR | |
| | | | Mean | ± | Mean | ± | Mean | ± | Mean | ± | Mean | ± | Mean | ± | Mean | ± |
| AC | 6 | Pre | 0.94 | 0.03 | 0.17 | 0.04 | 0.88 | 0.03 | 0.93 | 0.03 | 0.18 | 0.06 | 0.87 | 0.04 | 13.42 | 5.87 |
| | | MUS | 0.92 | 0.04 | 0.15 | 0.01 | 0.88 | 0.03 | 0.26 | 0.13 | 0.29 | 0.07 | 0.49 | 0.04 | 148.08 | 23.38 |
| | | PBS | 0.98 | 0.01 | 0.14 | 0.04 | 0.92 | 0.02 | 0.94 | 0.04 | 0.21 | 0.08 | 0.86 | 0.04 | 13.08 | 5.70 |
| IC | 7 | Pre | 0.88 | 0.05 | 0.23 | 0.05 | 0.83 | 0.04 | 0.87 | 0.04 | 0.32 | 0.06 | 0.77 | 0.03 | 16.71 | 6.03 |
| | | MUS | 0.40 | 0.13 | 0.13 | 0.04 | 0.64 | 0.05 | 0.03 | 0.01 | 0.06 | 0.04 | 0.48 | 0.02 | 187.29 | 12.71 |
| | | PBS | 0.93 | 0.03 | 0.21 | 0.03 | 0.86 | 0.02 | 0.76 | 0.12 | 0.30 | 0.05 | 0.73 | 0.05 | 29.50 | 10.42 |
| PPC | 7 | Pre | 0.99 | 0.01 | 0.13 | 0.03 | 0.93 | 0.01 | 0.96 | 0.01 | 0.32 | 0.08 | 0.82 | 0.04 | 28.93 | 9.08 |
| | | MUS | 0.94 | 0.03 | 0.35 | 0.10 | 0.80 | 0.06 | 0.31 | 0.34 | 0.20 | 0.03 | 0.55 | 0.04 | 133.21 | 32.37 |
| | | PBS | 0.96 | 0.02 | 0.24 | 0.03 | 0.86 | 0.01 | 0.90 | 0.05 | 0.32 | 0.09 | 0.79 | 0.05 | 42.29 | 12.55 |

| Target | mouse # | # of neurons | R#1 Hit rate | R#1 FA rate | R#1 Correct rate | R#2 Hit rate | R#2 FA rate | R#2 Correct rate | TTR |
|---|---|---|---|---|---|---|---|---|---|
| AC | 602 | 40 | 1.00 | 0.11 | 0.95 | 0.95 | 0.06 | 0.95 | 21 |
| | | 42 | 0.66 | 0.16 | 0.75 | 1.00 | 0.03 | 0.99 | 41 |
| | | 56 | 0.68 | 0.01 | 0.84 | 0.91 | 0.02 | 0.95 | 6 |
| | 605 | 37 | 0.96 | 0.16 | 0.90 | 0.97 | 0.01 | 0.98 | 16 |
| | | 89 | 0.93 | 0.17 | 0.88 | 0.80 | 0.31 | 0.75 | 66 |
| | | 64 | 0.42 | 0.01 | 0.71 | 0.77 | 0.11 | 0.83 | 18 |
| | 606 | 37 | 0.91 | 0.18 | 0.86 | 0.86 | 0.15 | 0.86 | 14 |
| | | 35 | 0.69 | 0.25 | 0.72 | 0.97 | 0.07 | 0.95 | 13 |
| AC total | N=3 | 400 | 0.78 ± 0.03 | 0.17 ± 0.04 | 0.83 ± 0.03 | 0.90 ± 0.03 | 0.10 ± 0.04 | 0.90 ± 0.03 | 24.38 ± 6.95 |
| IC | 602 | 22 | 0.93 | 0.26 | 0.84 | 0.95 | 0.42 | 0.77 | 45 |
| | | 58 | 0.83 | 0.41 | 0.71 | 1.00 | 0.14 | 0.93 | 63 |
| | | 13 | 0.97 | 0.41 | 0.78 | 0.98 | 0.12 | 0.93 | 22 |
| | 606 | 9 | 0.96 | 0.07 | 0.95 | 0.96 | 0.05 | 0.96 | 51 |
| | | 20 | 1.00 | 0.22 | 0.89 | 0.85 | 0.14 | 0.86 | 22 |
| | | 23 | 1.00 | 0.19 | 0.91 | 0.96 | 0.03 | 0.97 | 5 |
| | | 24 | 1.00 | 0.24 | 0.88 | 0.97 | 0.04 | 0.97 | 22 |
| | 609 | 10 | 0.86 | 0.44 | 0.71 | 0.96 | 0.10 | 0.93 | 88 |
| IC total | N=3 | 179 | 0.94 ± 0.02 | 0.28 ± 0.05 | 0.83 ± 0.03 | 0.95 ± 0.02 | 0.13 ± 0.04 | 0.91 ± 0.02 | 39.75 ± 9.61 |

| Group | # of mice | Type | Condition | R1 | | | | | | R2 | | | | | | | |
|---|---|---|---|---|---|---|---|---|---|---|---|---|---|---|---|---|---|
| | | | | Hit rate | | FA rate | | Correct rate | | Hit rate | | FA rate | | Correct rate | | TTR | |
| | | | | Mean | ± | Mean | ± | Mean | ± | Mean | ± | Mean | ± | Mean | ± | Mean | ± |
| $PPC_{AC}$ | 14 | Type I | Light on | 0.96 | 0.02 | 0.22 | 0.05 | 0.87 | 0.02 | 0.74 | 0.06 | 0.37 | 0.04 | 0.68 | 0.03 | 66.79 | 10.98 |
| | | | Light off | 0.98 | 0.01 | 0.25 | 0.05 | 0.86 | 0.02 | 0.88 | 0.03 | 0.28 | 0.05 | 0.80 | 0.02 | 31.29 | 5.37 |
| | 6 | Type II | Light on | 1.00 | 0.04 | 0.12 | 0.04 | 0.94 | 0.02 | 0.92 | 0.04 | 0.34 | 0.08 | 0.79 | 0.03 | 23.00 | 6.84 |
| | | | Light off | 1.00 | 0.00 | 0.15 | 0.05 | 0.92 | 0.02 | 0.90 | 0.05 | 0.31 | 0.06 | 0.80 | 0.02 | | |
| $PPC_{IC}$ | 14 | Type I | Light on | 0.95 | 0.02 | 0.24 | 0.05 | 0.85 | 0.03 | 0.51 | 0.08 | 0.21 | 0.04 | 0.65 | 0.04 | 104.08 | 14.95 |
| | | | Light off | 0.97 | 0.01 | 0.22 | 0.04 | 0.88 | 0.02 | 0.91 | 0.03 | 0.29 | 0.03 | 0.81 | 0.02 | 24.57 | 5.09 |
| | 5 | Type II | Light on | 0.95 | 0.04 | 0.40 | 0.11 | 0.77 | 0.07 | 0.94 | 0.02 | 0.46 | 0.15 | 0.74 | 0.08 | 12.40 | 2.14 |
| | | | Light off | 0.98 | 0.01 | 0.30 | 0.07 | 0.84 | 0.04 | 0.95 | 0.03 | 0.21 | 0.08 | 0.87 | 0.04 | | |
| $PPC_{ctrl}$ | 6 | Type I | Light on | 0.98 | 0.01 | 0.29 | 0.07 | 0.85 | 0.04 | 0.93 | 0.03 | 0.29 | 0.05 | 0.82 | 0.04 | 18.83 | 2.77 |
| | | | Light off | 0.98 | 0.02 | 0.19 | 0.06 | 0.89 | 0.03 | 0.93 | 0.03 | 0.27 | 0.05 | 0.83 | 0.03 | 24.00 | 3.89 |

| | | R1 | | | TR | | | R2 | | | |
|---|---|---|---|---|---|---|---|---|---|---|---|
| Group | # of neurons | Hit rate | FA rate | Correct rate | Hit rate | FA rate | Correct rate | Hit rate | FA rate | Correct rate | TTR |
| $PPC_{AC}$ | 149 | 0.99 | 0.50 | 0.75 | 0.77 | 0.92 | 0.42 | 0.97 | 0.39 | 0.79 | 27 |
| | 180 | 0.98 | 0.18 | 0.90 | 0.60 | 0.50 | 0.55 | 0.98 | 0.48 | 0.75 | 12 |
| | 92 | 1.00 | 0.27 | 0.87 | 0.85 | 0.82 | 0.50 | 0.84 | 0.18 | 0.83 | 43 |
| | 54 | 0.80 | 0.38 | 0.71 | 0.57 | 0.58 | 0.49 | 0.74 | 0.18 | 0.78 | 74 |
| | 72 | 0.97 | 0.66 | 0.66 | 1.00 | 0.81 | 0.58 | 0.98 | 0.29 | 0.85 | 32 |
| | 47 | 0.98 | 0.22 | 0.88 | 0.70 | 0.88 | 0.44 | 0.92 | 0.28 | 0.82 | 19 |
| $PPC_{AC}$ total | 594 | 0.95 ± 0.03 | 0.37 ± 0.08 | 0.79 ± 0.04 | 0.75 ± 0.07 | 0.75 ± 0.07 | 0.50 ± 0.02 | 0.90 ± 0.04 | 0.30 ± 0.05 | 0.80 ± 0.01 | 34.5 ± 9.02 |
| $PPC_{IC}$ | 57 | 1.00 | 0.38 | 0.81 | 0.80 | 0.75 | 0.50 | 0.87 | 0.14 | 0.86 | 23 |
| | 37 | 1.00 | 0.78 | 0.61 | 0.73 | 0.82 | 0.45 | 1.00 | 0.22 | 0.89 | 23 |
| | 5 | 1.00 | 0.39 | 0.81 | 1.00 | 0.67 | 0.60 | 0.98 | 0.19 | 0.90 | 6 |
| | 4 | 0.99 | 0.09 | 0.95 | 0.83 | 0.67 | 0.58 | 0.95 | 0.29 | 0.83 | 13 |
| | 1 | 1.00 | 0.18 | 0.91 | 0.83 | 0.75 | 0.54 | 0.92 | 0.32 | 0.80 | 25 |
| | 13 | 0.61 | 0.65 | 0.48 | 0.50 | 0.64 | 0.43 | 0.89 | 0.51 | 0.69 | 29 |
| | 14 | 0.76 | 0.24 | 0.76 | 1.00 | 0.00 | 1.00 | 0.82 | 0.20 | 0.81 | 5 |
| | 10 | 0.94 | 0.41 | 0.77 | 0.89 | 0.71 | 0.59 | 0.78 | 0.07 | 0.85 | 57 |
| | 27 | 0.99 | 0.34 | 0.83 | 0.50 | 0.70 | 0.39 | 0.79 | 0.50 | 0.65 | 19 |
| | 11 | 0.35 | 0.17 | 0.59 | 0.36 | 0.30 | 0.52 | 0.76 | 0.10 | 0.83 | 22 |
| | 10 | 0.88 | 0.08 | 0.90 | 0.67 | 0.60 | 0.55 | 0.82 | 0.18 | 0.82 | 12 |
| | 5 | 0.93 | 0.43 | 0.75 | 0.17 | 0.23 | 0.47 | 0.75 | 0.22 | 0.76 | 61 |
| $PPC_{IC}$ total | 194 | 0.87 ± 0.06 | 0.34 ± 0.06 | 0.76 ± 0.04 | 0.69 ± 0.07 | 0.57 ± 0.07 | 0.55 ± 0.05 | 0.86 ± 0.02 | 0.25 ± 0.04 | 0.81 ± 0.02 | 24.58 ± 5.12 |
